# Acquisition of non-olfactory encoding improves odour discrimination in olfactory cortex

**DOI:** 10.1101/2023.07.04.547685

**Authors:** Noel Federman, Sebastián A. Romano, Macarena Amigo-Duran, Lucca Salomon, Antonia Marin-Burgin

**Affiliations:** Instituto de Investigación en Biomedicina de Buenos Aires (IBioBA)-CONICET-Partner Institute of the Max Planck Society; Godoy Cruz 2390, C1425FQD, Buenos Aires, Argentina

## Abstract

Primary sensory cortices, initially considered elementary encoders of physicochemical attributes of environmental stimuli, are now known to be modulated by other aspects of experience, such as attentional state and internal expectations ^1–3^, movement-related signals ^4–7^ and spatial information ^2, 8, 9^. However, the specific role of these signals in cortical sensory processing is not fully understood ^10^. Here we reveal multiple and diverse non-olfactory responses in the primary olfactory (piriform) cortex (PCx), which dynamically enhance PCx odour discrimination according to behavioural demands. We designed a behavioural task using a virtual reality environment and performed recordings in PCx neurons. In this task, mice were trained to associate specific odours with visual contexts in order to receive a reward. We found that learning shifts PCx activity from encoding solely odour identity to a more complex regime. In this regime, positional, contextual, and associative responses emerge on odour-responsive neurons that thus become mixed-selective. Contextual information is sustained in PCx activity of expert animals, specifically when visual context identity is needed to solve the task. After learning, odours are better decoded from PCx activity when mice are engaged in the task and when odours are presented within a rewarded context. This enhancement of PCx olfactory processing is reliant on the acquired mixed-selectivity. Thus, the integration of extra-sensory inputs within primary sensory cortices can encode the behavioural relevance of encountered stimuli while improving sensory processing.

## Main

Stimulus-evoked activity in primary sensory cortices (PSCs) is thought to determine the identity of modality-specific stimuli. Nevertheless, recent studies have shown prominent non-sensory responses in these brain regions ^3–9^, but the functional role that these diverse modulations may play in sensory processing is still under investigation.

The ability to integrate sensory, behavioural and task-related information at the initial stages of cortical sensory processing could be advantageous: rather than simply relaying segregated information to higher-order association brain areas for subsequent processing, early integration at PSCs could enable state-and context-dependent sensory representations, allowing animals to flexibly process sensory information according to their current behavioural demands ^11^. The signature of such integration at the single-neuron level would be the presence of mixed-selectivity neurons that simultaneously encode multiple task variables of diverse nature. Mixed-selectivity is typically observed at higher order regions like the prefrontal cortex ^12–14^ and hippocampus ^15^, but limited research has focused on investigating mixed-selectivity in PSCs ^16^.

Here, we focused on the piriform cortex (PCx), the largest region of the mouse primary olfactory cortex that collects odour information from the olfactory bulb to encode odour identity ^17–19^ and also receives inputs from higher order processing areas ^20–24^, suggesting that it can be modulated by other aspects of sensory experience. Recently, it was shown that the posterior PCx can encode the spatial location of odour stimuli ^8^. How this non-olfactory information influence olfactory processing is not understood.

## Mice learn to discriminate the same odour in different spatial contexts

We developed a behavioural task in which mice have to learn that a particular odour is rewarded when presented at the entry of a specific visual context (Fig. 1a, Extended data Fig. 1a, b). Thirsty animals were trained to traverse a virtual linear corridor in order to reach a spatial location where one of two distinct visual contexts could be presented (green or grey). When entering the context, they received a 1-second-long puff of one of two possible odours (isoamyl acetate or ethyl butyrate). When reaching the reward zone at context exit, mice could choose to lick (GO response) or not to lick (NO-GO response) a reward spout to trigger the delivery of a water drop. Only one of the four possible odour-context combinations would lead to water delivery (rewarded odour-context; O_R_C_R_; Fig. 1b). Mice had to learn to discriminate the target O_R_C_R_ association with GO responses only on those trials. Importantly, the behavioural task was designed so that while mice run through the virtual corridor, they were able to see the approaching visual context in anticipation of odour delivery (Fig.1a and Extended data Fig. 1b). This allowed us to evaluate how contextual and olfactory information are individually related to behaviour and neuronal activity.

**Fig. 1.**
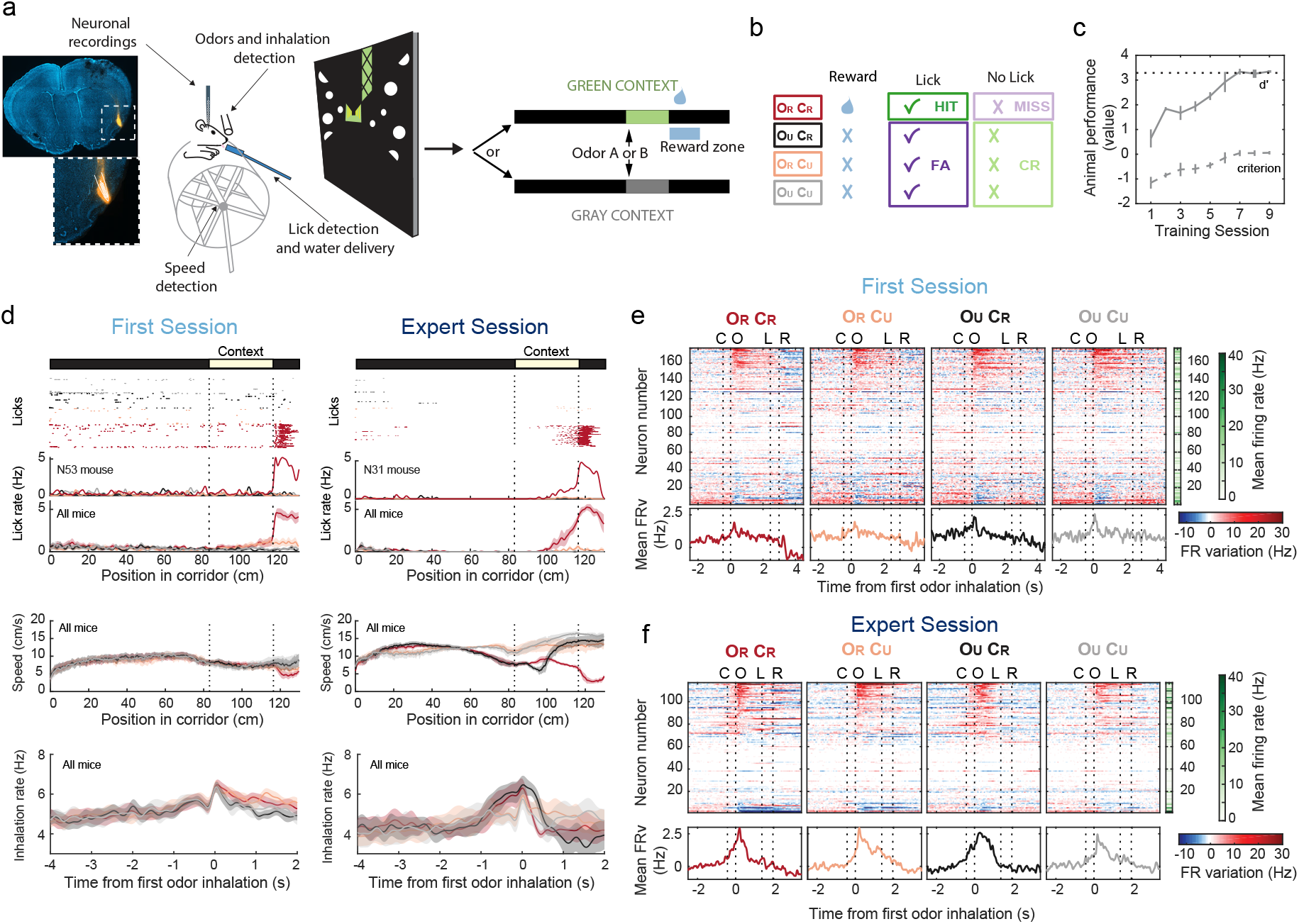
Learning an associative odour-virtual context-reward task. **a**, Schematic of the virtual reality setup. Example of a recording site with the position of the silicon probe stained with dye I. Mice can run through a corridor with two alternative visual contexts and odours. Delivery of odours occurs after entering visual context and the reward delivery occurs at the exit. **b,** Only one combination of context and odour is associated with the water reward. OR and OU, Odour rewarded and unrewarded. CR and CU, visual context rewarded and unrewarded. Depending on context, red/pink or black/grey represent different odours. CR, correct rejection, FA, false alarm. **c,** Behavioural performance (behavioural d’ and criterion, see Materials and Methods). Dashed line corresponds to 95% hits with 5% FA responses. n=4-10 mice. **d**, Changes in behavioural variables through learning. Top rasters show licks (dots) aligned to position in the corridor of two example animals (first session and expert session) where trial type is colour coded as in b. Lower panels show average licking rate, speed and inhalation rate on different trial types for first session (n=6 mice) and expert animals (n=4 mice). **e-f,** Colour maps showing all neuronal responses as variation in firing rate (FR; variation around the neuron’s mean FR) in first session (e) and expert session (f) animals aligned to the time of the first odour inhalation. Vertical dashed lines indicate alignments to median times of the following events: C: Context entry, O: odour delivery, L: licking response before water delivery, R: reward delivery. FR stitched for visualization (see Materials and Methods). Green color-coded column on the right shows the mean FR of each neuron. Neurons were sorted according to their average responses to ORCR trials. Lower panels show the mean FR variation (FRv) across the neuronal population for each trial type.

Animals became experts after around 7 days of training (Fig. 1c). Several task-related behavioural variables such as locomotion speed and inhalation rate were adjusted with learning, and this adjustment depended on animal position and sensory stimuli conditions (Fig. 1d). Moreover, learning induced anticipatory behaviours, such as an increase in sniffing rate before odour delivery and anticipatory licking responses when entering the rewarded zone (Fig. 1d), indicating that expert animals use odour in combination with visual contexts but also corridor position to guide their behaviour.

## Population dynamics in piriform cortex reveals spatial and associational signals that emerge with learning

To investigate whether PCx neural responses change as animals learned to associate odours with visual contexts and rewards, we performed acute recordings of PCx neuronal activity of animals in their first training session (177 neurons from 6 animals, 29.5 +/- 21.1 neurons per animal; mean +/- s.d) and animals that reached expert performance (117 neurons from 4 animals, 29.2 +/- 23.6 neurons per animal; mean +/- s.d.; Fig. 1a, e, f). There were no significant differences in the firing rates recorded in both conditions (Extended data Fig. 1c).

To analyse if we could extract odour information from these recordings at the population level, we cross-correlated the activity between trial types in first-session and expert animals. We pooled recordings and calculated a time series of population activity vectors containing the spike counts of each cell, computed in a sequence of time bins around odour stimulation. The similarity between these vectors evoked by different trial types and at different times during first sessions and expert sessions was quantified by the Pearson correlation coefficient. First-session evoked activity patterns following odorant stimulation that distinguished odour identity (Fig. 2a, two diagonal blocks in top panel). Analysis of activity patterns from expert animals (Fig. 2a, bottom panel) points to a more complex scenario: trial types can be initially grouped by visual context as animals approach the context entry (Fig. 2a, checkerboard pattern emerging before odorant stimulation in expert animals), later response patterns discriminate between odours (Fig. 2a, diagonal blocks around 0.3s in expert animals), and by 1.2s after first odour inhalation they fully differentiate rewarded trials from the rest of conditions (Fig. 2a, O_R_C_R_ trials in expert animals). We obtained a comparable result through principal component analysis (PCA). Projecting the time series of evoked activity patterns from expert recordings in principal component space resulted in trajectories that diverge first according to contexts, then to odours, finally leading to a deviation of rewarded trials trajectory (Fig. 2b). These results indicate that non-olfactory signals, presumable related to visual context and reward, can be observed in the activity of the population of PCx neurons. Interestingly, those responses were acquired with learning.

**Fig. 2.**
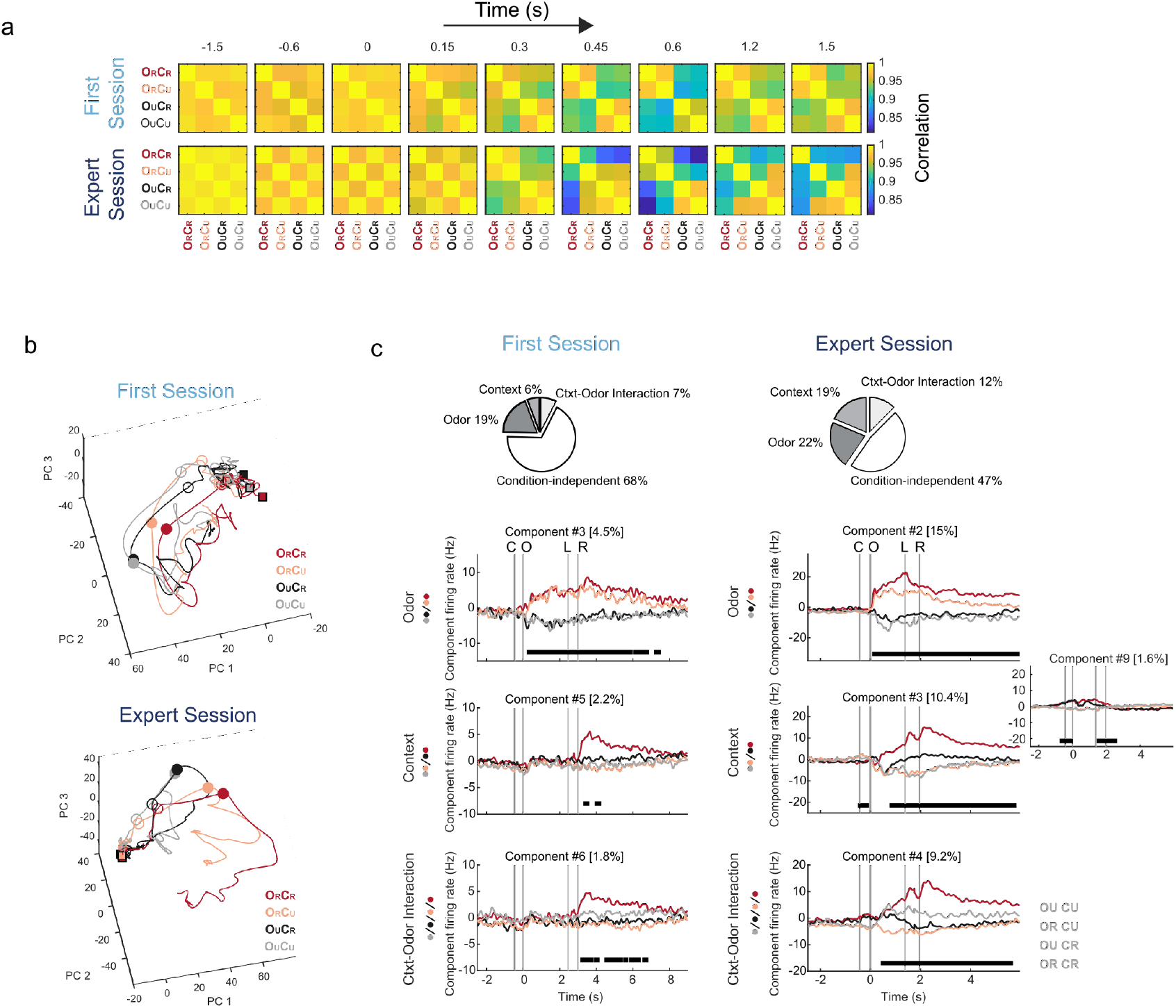
Population dynamics in the piriform cortex reveals spatial and associational signals. **a**, Pearson correlation coefficients of the time series of population activity vectors containing the spike counts of each cell, computed in a sequence of time bins around odour stimulation, for each trial type. **b,** Principal component analysis (PCA) of the pooled time series of evoked activity patterns for first-session and expert-session recordings. Filled squares, empty circles and filled circles label -2.5 seconds, 0 seconds and 0.2 seconds relative to first odour inhalation, respectively. Colours indicate trial type. **c,** Demixed principal components (dPCA). Pie charts show how the total signal variance is split among task parameters. Top panels: first odour discrimination components, middle panels: first context discrimination components (inset in expert session also shows the second component), bottom panels: first context-odour interaction component. First-session recordings on the left, and expert-session recordings on the right. In each subplot, the full data are projected onto the respective dPCA decoder axis. Thick black lines show time intervals during which the respective task parameters (odour, context, and context-odour interaction) can be significantly decoded from single-trial activity (see Materials and Methods). Variances explained by each component are shown as percentages. More dPCA components are shown in Extended data Fig. 2.

To quantify which of the task-related variables significantly contribute to PCx neuronal activity, we decomposed the time-varying contributions of sensory stimuli conditions to the population activity dynamics by applying demixed PCA (dPCA;^25^) across odour-context combinations (Fig. 2c, Extended data Fig. 2a, c; *Materials and Methods*). dPCA takes into consideration the different task conditions to infer components that depend on single parameters, allowing to disentangle data dependencies on specific variables, like odours, contexts and reward. In both first session and expert animals, signal variance had strong contributions from odour components (Fig. 2c, pie charts) and odour information can be decoded from several components that show either transient or sustained activity (Fig. 2c and Extended data Fig. 2b, d). Contributions from contextual and context-odour interaction components were larger in expert animals (Fig. 2c, pie charts), and contextual information could be decoded from components that anticipate the entry and exit of the visual context zone (Fig. 2c and Extended data Fig. 2b, d). Reward information is observed as a peak of contextual and context-odour interaction components in O_R_C_R_ trials after reward consumption in both first-session and expert animals. Nevertheless, in expert animals, some components could discriminate O_R_C_R_ trials before the animal’s decision to lick for reward (Fig. 2c and Extended data Fig. 2d, see components #3, 4 and 12 in Experts). These observations indicate that, only after learning, population PCx activity encodes information about spatial and associational aspects of the task. How are these population representations composed?

## Learning induces mixed-selectivity encoding in single neurons of piriform cortex

The observed olfactory and contextual signals could be conveyed by different neuronal sub-populations of PCx neurons. Alternatively, this information could be merged by single neurons. We evaluated this by studying single PCx neuronal responses from first-session and expert animals performing the task (Fig. 3a). As expected by previous work ^17, 26, 27^, PCx neurons showed typical excitatory and inhibitory responses to odorant identity locked to odour inhalation onset, and displayed respiratory-locked activity (Fig. 1e-f, Fig. 3b, c and Extended data Fig. 3d, h). We also found neurons whose activity levels varied according to the animal’s running speed, along with changes in firing rate preceding and/or following single licking events across all trial types and following reward consumption in O_R_C_R_ trials (Fig. 3b, c and Extended data Fig. 3b, c, f, g). Interestingly, once animals learned the task, we found neurons that fired according to the animal spatial location along the virtual corridor (Fig. 3c, “Position responses”) and neurons that displayed a diversity of discriminative responses to the different odour-context combinations (Fig. 3c and Extended data Fig. 3i, “Associative responses”). Some of these neurons were selective for the rewarded O_R_C_R_ trials following odorant stimulation but in anticipation of the animal’s licking response (Fig. 3c and Extended data Fig. 3i). These results show that beyond odour identity encoding, the activity of individual neurons in PCx is rich in behavioural information, capable of context-dependent olfactory processing and displays learning-related associative and choice signals.

**Fig. 3.**
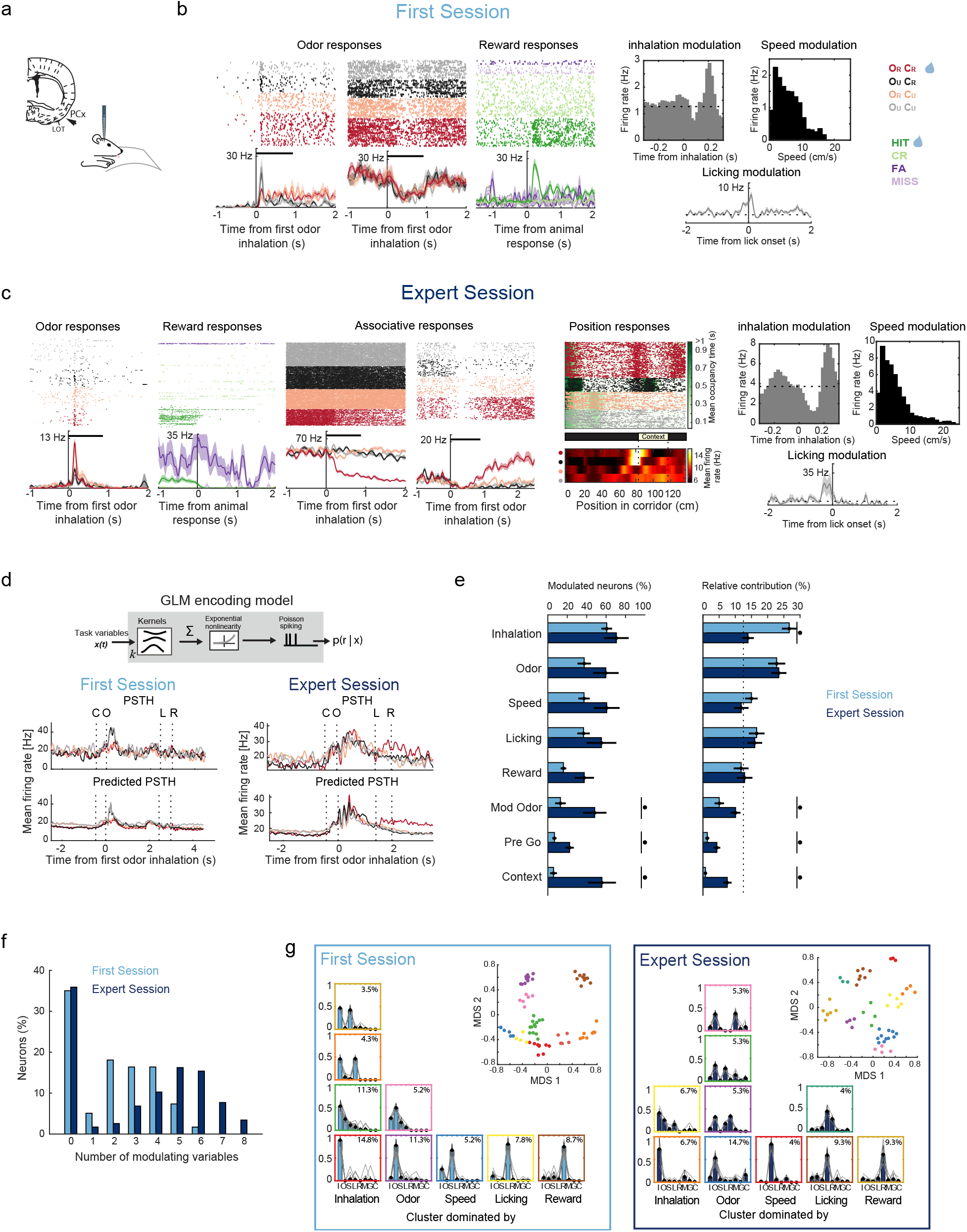
Single-cell responses across learning. **a,** Schematic of PCx recording site in behaving animals. **b,** Examples of neurons from mice recorded during first sessions. Odour and reward responses are shown in raster plots of action potentials (ticks) colour coded by trial type (odour responses) or trial outcome (reward responses), according to the colour code on the right. Odour pulse duration is indicated with a horizontal black bar. Bottom panels show the average firing rate of the neurons. Modulation by inhalation, speed and licking are also shown in the right three panels. **c**, Examples of neurons from mice recorded during expert sessions. Odour, reward, inhalation, speed and licking responses are shown as in B. Expert animals also show associative responsive neurons as well as neurons responding to the animal position along the corridor. Position responses in the bottom panel are normalized by the time the animal spent in each location (animal’s mean occupancy time is shown in green under the raster plot). **d**, Top, scheme of the generalized linear model (GLM) applied to single-neuron single-trial recordings to extract linear kernels that specify the neuron’s dependence on every task variable. Bottom, examples of peri-stimulus time histograms (PSTH) and model-predictions of PSTHs of one neuron recorded during a first session (top panels) and one during an expert session (bottom panels). Recordings were aligned according to median times of the following trial events: C: Context entry, O: odour delivery, L: licking response before water delivery, R: reward delivery. PSTHs stitched for visualization (see Materials and Methods). **e,** Percentage of modulated neurons for each task variable (left) and the relative contribution of each task variable to this modulation (right; dashed line indicates equal contribution by all variables). Black dots indicate statistically significant differences between first or expert session recordings (Wilcoxon rank sum test. Left panel: ModOdour, p-value = 3.8 x 10-2; preGo, p-value = 1.9 x 10-2; Context, p-value = 9.5 x 10-3. Right panel: Inhalation, p-value = 1.4 x 10-4; ModOdour, p-value = 2 x 10-7; preGo, p-value = 1.9 x 10-4, Context, p-value = 1.5 x 10-17). Light blue for first session neurons, dark blue for expert session neurons. **f,** Percentage of neurons modulated by different total number of variables. The number of variables modulating PCx neuronal activity increased with learning (Wilcoxon rank sum test for tied data; p=1.1 x 10-4). **g**, Clustering neurons according to the relative contribution of their encoded task variables. Each coloured square represents the mean relative contribution of the variables of individual clusters (mean +/- std). Grey lines show individual neurons in the cluster. Percentages indicate the proportion of neurons included in each cluster relative to the total population. Columns of bar plots are organized according to the dominating variable (labelled below each column, I: inhalation, O: odour, S: speed, L: licking, R: reward, M: modulation of odour by context, G: Pre Go modulation, C: context). Top-right inset shows the multidimensional scaling (MDS) projection of the relative contribution of the encoded variables. Each dot is a neuron, and neurons with similar relative contributions occupy similar regions of the MDS plane. Neurons are colour-coded according to the cluster to which they belong (same colour-codes of bar plots).

To characterize the mixture of sensory and behavioural signals that simultaneously affect the activity of single neurons, we implemented a statistical approach based on a Poisson generalized linear model (GLM) of neuronal encoding ^28, 29^ (Fig. 3d, Extended data Fig. 4a, b; *Materials and Methods*). The model fits single-trial time-varying spike responses against the timing and value of each task variable during each trial. (Extended data Fig. 4a). For each neuron, we selected an encoding model containing the combination of these variables that maximized model performance in predicting neuronal activity from trials held out from the fitting procedure (Extended data Fig. 4c). In doing so, it infers linear filters (or kernels) that quantify the dependencies of neuronal responses on each variable (Fig. 3e, f, Extended data Fig. 5a, b, Extended data Fig. 6).

The variables considered in our model were sensory, motor and cognitive: rewarded and unrewarded odours (O_R_ and O_U_); animal position along rewarded and unrewarded visual contexts (C_R_ and C_U_); reward consumption (R); inhalation (I); running speed (S); licking (L); modulation of odour responses by presence of rewarded context (modO_R_ and modO_U_); anticipation to GO response (preGO). For each neuron, we selected an encoding model containing the combination of these variables that maximized model performance. Model predictions of peri-stimulus time histograms (PSTHs) aligned to task events (Extended data Fig. 6) explained a larger degree of variance of pooled expert-session data (66%) compared to pooled first-session data (24%), even after matching the total number of trials between both conditions (Experts: 49%, First-session: 23%). This is in line with the previous observation that individual stimulus and contextual dPCA components better capture variance in population data from expert-session recordings than from first session (Extended data Fig. 2 a, c). Together, these results point to a stronger and more reliable modulation of PCx neuronal activity by sensory and behavioural variables after learning.

The fitted models reveal that single PCx neurons carry information about a variety of olfactory and non-olfactory variables in distinct ways, differing in magnitude and temporal dynamics (Extended data Fig. 5 and Extended data Fig. 6). The neuronal activity of first-session animals was dominated by inhalation and odour kernels (Fig. 3e). At the level of the fitted kernels, learning is accompanied by two prominent features: the presence of strong modO_R_ and modO_U_ kernels that modulate odour responses in a context dependent manner a few hundred milliseconds following olfactory stimulation, and the incorporation of C_R_ and C_U_ kernels that encode the entry and exit cues of the visual contexts (Fig. 3e and Extended data Fig. 5c). Neurons from expert animals were modulated by a larger number of parameters. This increase in multiplexing after learning was confirmed by estimating the percentage of neurons modulated by each variable (Fig. 3f), the absolute and relative contribution of the variables to the model (Fig. 3e, right panel and Extended data Fig. 4d), and the amplitude of the obtained kernels (Extended data Fig. 4e and Extended data Fig. 5c). For all these quantities, expert-session neurons showed a larger extent of visual context encoding, modulations of odour responses by rewarded context, and activity in anticipation to GO responses. In addition, licking and reward consumption had larger absolute contributions in experts, while reward consumption also showed larger kernel amplitude (Extended data Fig. 4d, e). Thus, learning embeds piriform cortex neurons with mixed selectivity responses to different aspects of the behavioural task.

## Learned mixed-selectivity responses on PCx are structured

We investigated whether learning reorganizes PCx neurons in new functional groups. The mixed-selective neurons observed after learning could be selective to any given mixture of variables, with no particular preference. On the contrary, learning could promote specific patterns of mixed selectivity composed of particular combinations of variables that modulate together single-neuron responses. To address this, we used a hierarchical clustering approach to identify assemblies of neurons with similar modulation profiles (*Materials and Methods*). Clustering neurons according to the relative contributions of the variables they encode (Fig. 3g, Extended data Fig. 7) showed that first-session profiles were typically dominated by contributions from single variables (Fig. 3g, bottom row, 47.8% of fitted cells), or pairs of variables that included either inhalation or odour kernels (Fig. 3g, upper rows of “Inhalation” and “Odour” columns, 24.3% of fitted cells). Clusters of expert-session neurons were less dominated by inhalatory drive and 30.6% of the fitted neurons were grouped around odour kernels combined with kernels related to visual context, modulations of odour responses by rewarded context, reward consumption and licking (Fig. 3g, “odour” column). This result suggests that learning-induced mixed selectivity reshapes PCx neuronal populations by specifically integrating olfactory information with variables important for the behavioural task.

## Decoding of sensory information improves with mixed-selective neurons

We wonder whether the mixed-encoding has functional consequences, affecting the amount of sensory information carried by PCx responses. To evaluate this, we used the fitted GLM encoding models and the observed spiking activity of the PCx, to perform model-based trial decoding of odour, context and reward information (see *Materials and Methods*). We found that, while the trial-by-trial activity of neurons in expert animals allows decoding the identity of odours and of visual contexts, the activity of neurons in first session animals only permits to decode information about odours (Fig. 4a). Trials with reward consumption are successfully decoded in both conditions (Extended data Fig. 8c).

**Fig. 4.**
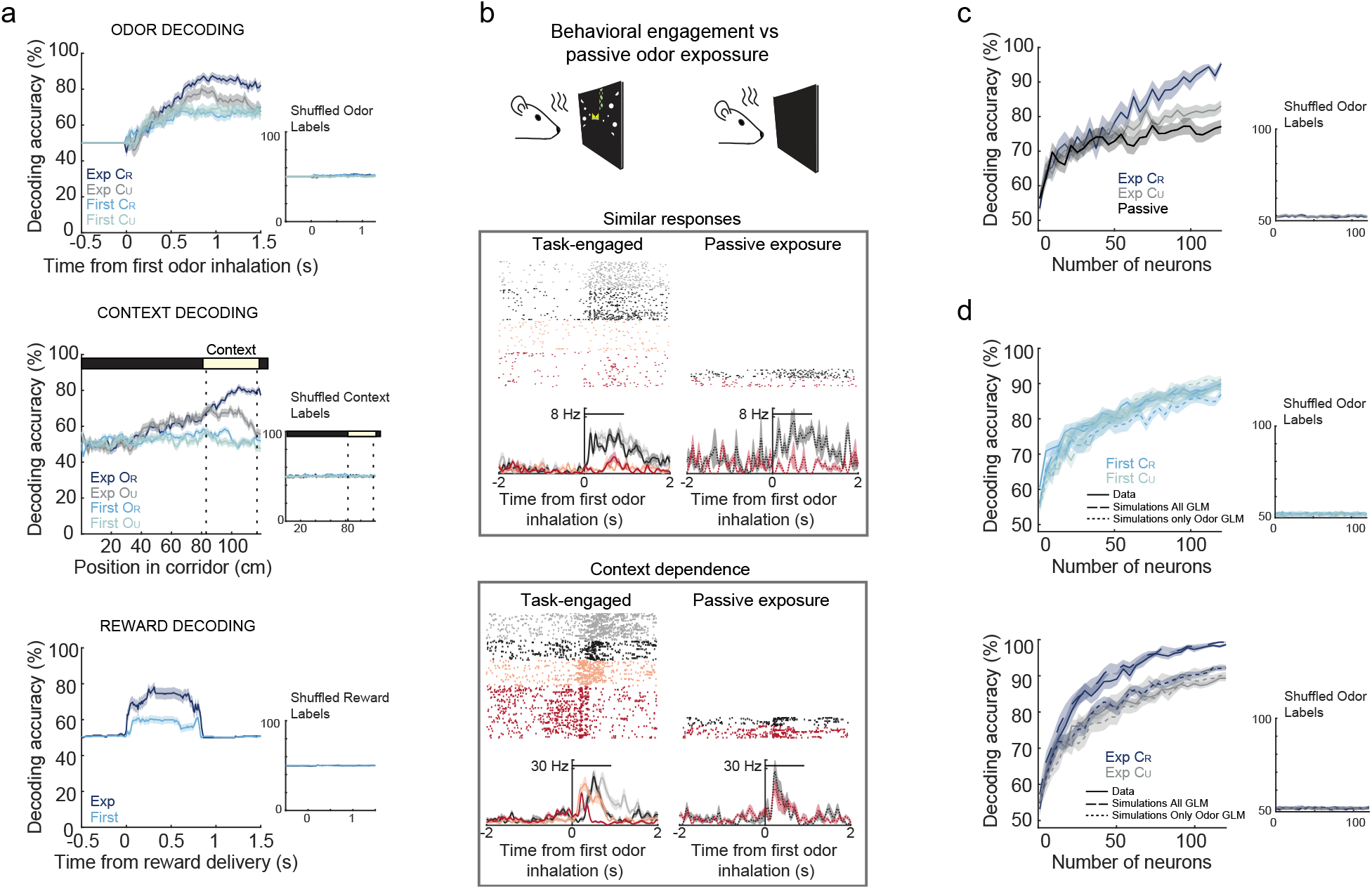
Learning induces multidimensional encoding in piriform neurons. **a,** GLM-based trial decoding of binary variable categories (rewarded vs unrewarded odour, rewarded vs unrewarded visual context, reward consumption vs no reward consumption), using first-session and expert-session recordings. The number of trials used for model fitting and decoding analysis was matched across different trial types. **b**, Top, the scheme represents the experimental condition, during task engagement or when the virtual reality was turn off and animals were passively exposed to odours. Bottom, examples of 2 neurons recorded in expert sessions during the task or under passive exposure to odours. The upper neuron has similar responses under both conditions, while the lower neurons show context-dependence of the response. **c**, Decoding accuracy of linear classifiers for odour identity as a function of neuronal population size, using expert-session data in CR and CU trials, and using neuronal responses from passive odour exposure. Decoding performed at the moment when accuracy first peaks (0.5 s and 1 s after first odour inhalation for task-engaged and passive conditions, respectively). Inset shows decoding accuracy after shuffling trial labels odour labels. **d**, Decoding accuracy of linear classifiers for odour identity as a function of neuronal population size, for recorded data (Data) and simulations of GLM models including all fitted kernels (Full GLM) and after removing contributions of non-odour-related kernels from the simulations of odour-responsive neurons (Only Odour GLM). Top, accuracy for first-session data and simulations in CR and CU trials. Bottom, same as Top but for expert-session. Insets show results after shuffling odour labels. Decoding always performed at 0.5 s after first odour inhalation. For all decoding analysis shown in a, c and d, the total number of trials used for training and testing classifiers was matched across the different trial types compared.

Notably, in expert animals, decoding of context in unrewarded odour trials decays to chance level once the odour was presented, while for the rewarded odour trials decoding performance is maintained at high levels (Fig. 4a). Thus, the learned contextual information is dynamically modulated along each trial, and persists in PCx only when context identity is needed to disambiguate reward outcome. This persistent activity is reminiscent of a working-memory trace that represents contextual information held in memory to determine trial performance, and further supports an associative function of PCx.

Although odour identity could be extracted from PCx activity of both expert and first session animals, odour decoding in expert animals reached a higher accuracy and needed a fewer number of neurons than before learning (Fig. 4a and Extended data Fig. 8a). Moreover, odour identity in expert animals was better decoded in trials when correct odour discrimination leads to a reward, that is, when the odour was presented in the rewarded visual context (Fig. 4a and Extended data Fig 8a). This indicates that the learned contextual modulation in the PCx enhances olfactory information during moments when odour discrimination becomes behaviourally significant.

If PCx non-olfactory modulation is indeed beneficial for odour decoding, we hypothesized that odour discrimination in PCx should be reduced when contextual modulation is absent, for example, when animals are passively exposed to the same odours. We recorded PCx neuronal responses during task engagement and, by subsequently turning off the virtual reality, the same neurons were recorded during passive exposure to both odours. Comparing neuronal activity under passive stimulation to that observed while performing the task revealed that some neurons have similar responses in both conditions, while others showed dramatic changes in their responses (Fig. 4b and Extended data Fig. 9). In particular, we found neurons with almost identical responses to both odours during passive stimulation, while the response of these neurons during task engagement allows discrimination of both odours and also of type of trial (Fig. 4b and Extended data Fig. 9). Using linear decoders on the recorded neuronal populations (see *Materials and Methods*), we observed a lower and slower performance of odour identity decoding under passive stimulation compared to the task-engaged condition (Fig 4c). These results suggest that at the population and even at the single neuronal level, the multimodal encoding acquired during learning leads to a faster and improved accuracy of the odour discrimination function of the PCx.

To specifically test if the acquired mixed-selectivity underlies the improvement of odour discrimination in rewarded contexts (Fig. 4a and Extended data Fig 8a), we used linear decoders on GLM simulations of PCx neuronal activity, which reproduced the odour decoding performance of the observed data (Fig. 4d). When we removed the contribution of non-olfactory kernels to the odour encoding neurons, odour decoding accuracy decreased in expert rewarded context trials, reproducing the decoding performance observed during the unrewarded context trials (Fig. 4d). We obtained similar results using GLM based decoding (Extended data Fig. 8d).

Interestingly, no single non-olfactory variable was responsible for the accuracy increase in rewarded context trials, and the effect was dependent on the presence of the multiple non-olfactory kernels of odour-coding neurons from expert animals (Extended data Fig. 8e). These findings demonstrate that learned multidimensional mixed-selectivity allows PCx neurons to enhance odour discrimination when odours acquire behavioural relevance.

## Discussion

In this work we examined how encoding in the primary olfactory cortex is modified when odours gain behavioural relevance through associative learning between odorants, visual contexts and rewards. We found that learning was associated with the emergence of widespread contextual non-olfactory modulations of PCx activity which, when integrated with its canonical olfactory responses, allow the PCx to better discriminate odours. The underlying computational mechanism is the acquisition of mixed-selectivity for diverse task-related variables by odour-responsive PCx neurons. This neuronal code is engaged in a dynamic manner depending on task demands, providing experience-dependent contextual and non-olfactory information to improve odour processing in the PCx.

Non-sensory modulations of primary sensory cortices had been found in other primary sensory cortices. Locomotion modulates activity in visual cortex V1 ^4–6^ and auditory cortex A1 ^30, 31^. Animal movements can modulate activity across the dorsal cortex, including V1 and somatosensory cortex S1 ^7, 32^. Spatial representation had been also found in V1 ^2, 9^ and more recently in PCx ^8^. Despite these remarkable results, the role of these diverse sources of non-sensory information in primary sensory cortices, and their influence in sensory processing in particular is still a matter of debate ^10, 33, 34^. Here we found that learning induces encoding of several task related variables that are dynamically used to improve odour decoding in the PCx. This is in line with the observation of the representation of odours in PCx drifts over the course of days ^35^, suggesting that the PCx is a fast and continual learning system.

Encoding based on neuronal “multitaskers” is often found in high-order prefrontal ^36, 37^, parietal^38^ and hippocampal ^15^ cortices. We show that learning shifts the mouse PCx cortex from the typical modality-specific sensory-driven functional organization of primary sensory cortices, towards a mixed-selectivity associational regime. This regime could facilitate separability by reducing overlapping odour representations in the PCx, promoting discrimination and avoiding generalization of the same odour in different contexts (Extended data Fig. 10). Furthermore, mixed-selectivity in the PCx could allow simple downstream decoding operations to produce a wide range of context-dependent decision-making responses, as has been suggested for association and executive brain regions ^14, 16, 39^.

Our findings provide insights into the plasticity of PCx encoding in response to experience and highlight a computational mechanism at a primary sensory cortex that exploits the multiplexed nature of sensory experience to improve neural coding. We speculate that a similar process could take place in other primary sensory cortices known to be modulated by non-sensory information.

---

**Extended data Fig. 1.**
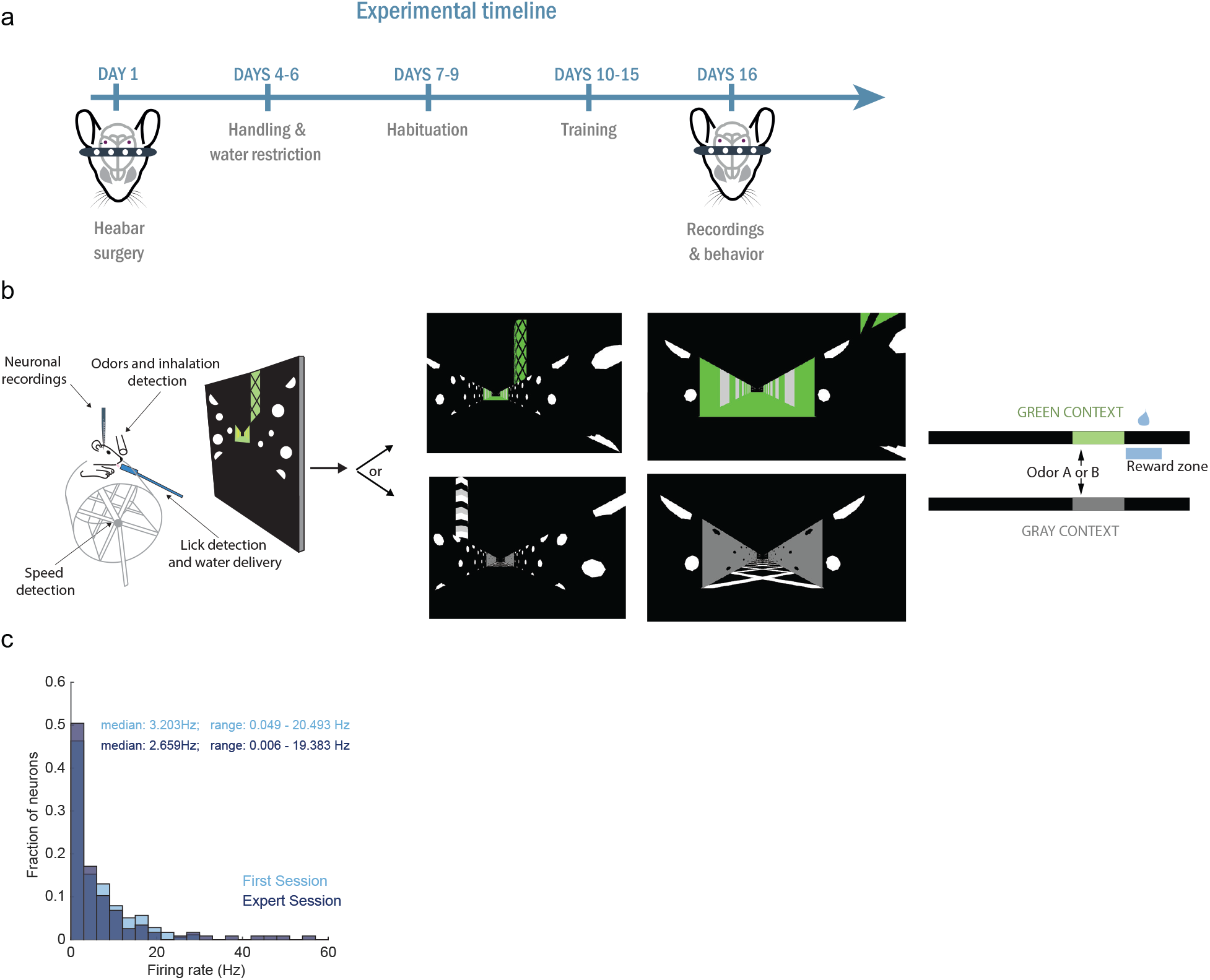
Experimental setup and timeline. **a**, Experiment timeline indicating the training and recording protocol. **b**, Experimental setup showing the two alternative virtual reality corridors (grey or green contexts) used in the experiments. **c,** Distribution of firing rates of recorded neurons for first-session recordings (light blue) and expert-session recordings (dark blue).

**Extended data Fig. 2.**
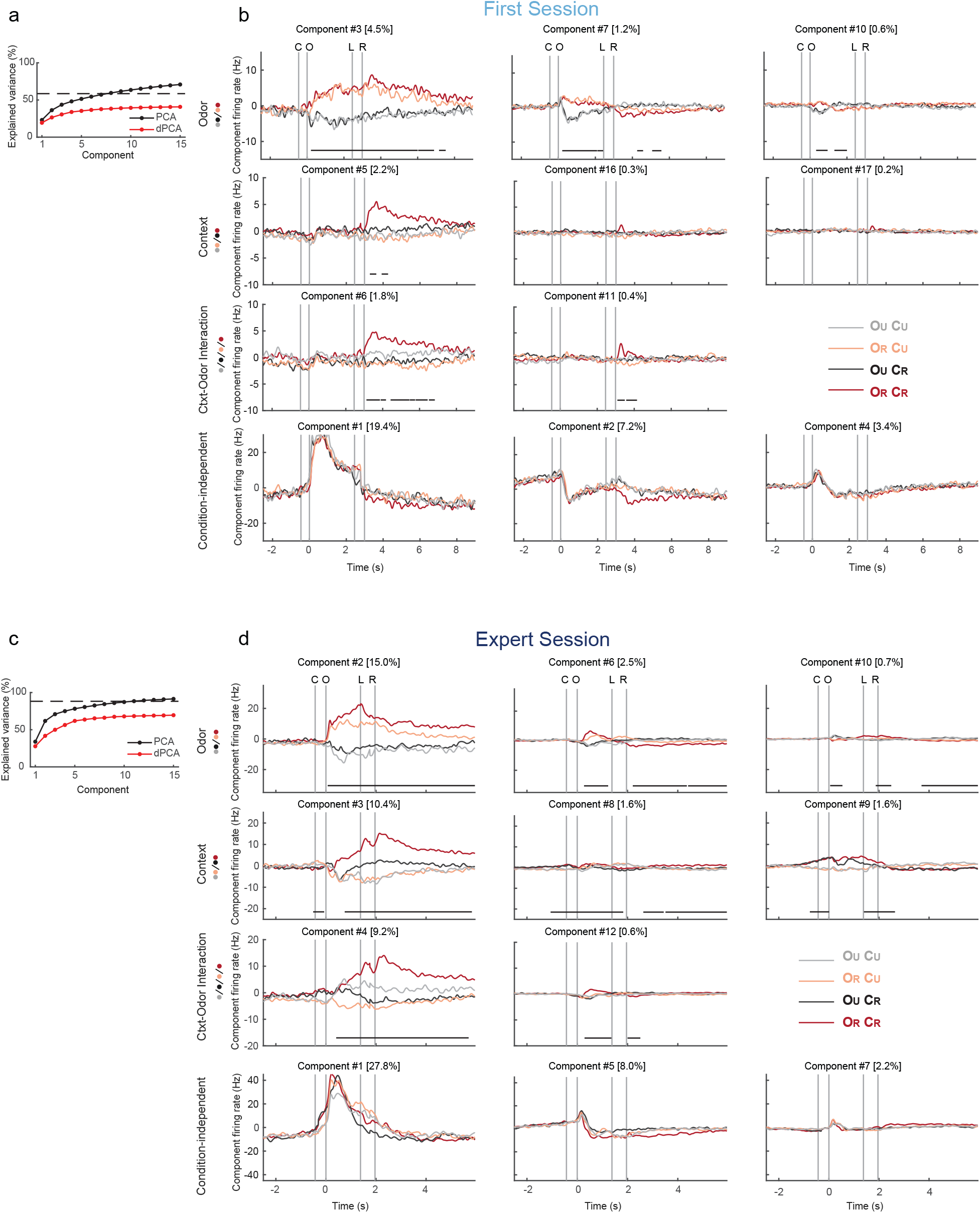
Demixed PCA components. **a and c,** Cumulative variance explained by dPCA and PCA from neurons recorded of animals in first session (a) or expert session (c). Dashed line shows an estimate of the fraction of signal variance in the data. **b and d,** Demixed principal components. From top to bottom panels: odour discrimination components, context discrimination components, context-odour interaction discrimination components, condition-independent components for first session (b) of expert session (d) recordings. In each subplot, the full data are projected onto the respective dPCA decoder axis. Thick black lines show time intervals during which the respective task parameters (odour, context, and context-odour interaction) can be significantly decoded from single-trial activity (see Materials and Methods).

**Extended data Fig. 3.**
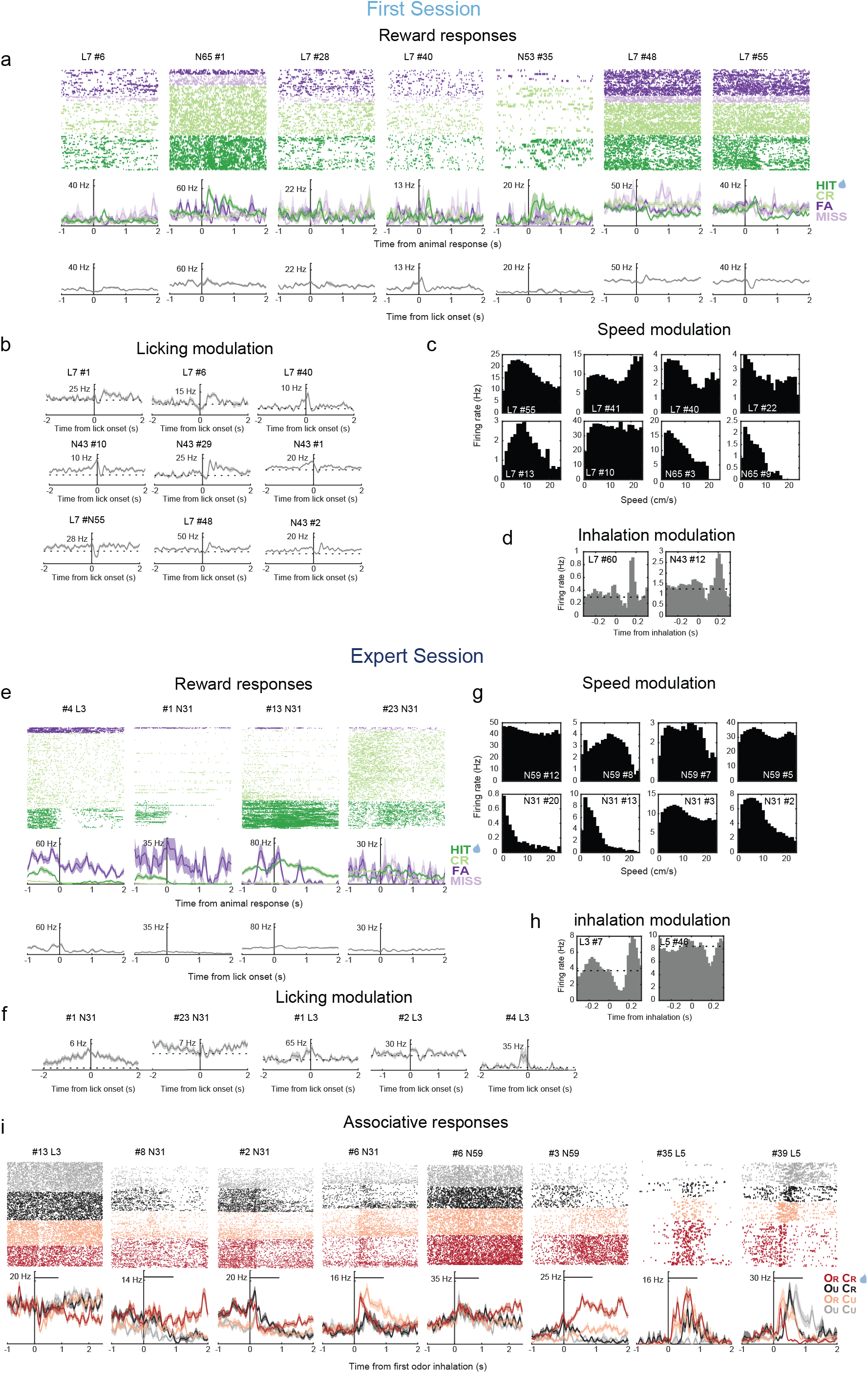
Examples of recorded PCx responses. **a and e,** Examples of reward responses are shown in raster plots of action potentials (ticks) colour coded by trial outcome (colour labels shown on the right) for first session (a) or expert sessions (e). Middle panels show the average firing rate of each raster aligned to the animal’s response after odour delivery. Bottom panels show the average firing rate of these neurons, aligned to the onset of every lick. **b and f,** Examples of responses of neurons modulated by licking in first sessions (b) or in expert sessions (f). Dashed line indicates mean firing rate. **c and g,** Examples of responses of neurons modulated by speed in first session (c) or in expert session (g). **d and h,** Examples of responses of neurons modulated by inhalation in first sessions (d) or in expert sessions (h). Dashed line indicates mean firing rate. (**I**) Examples of neurons with associative responses in expert animals. Horizontal black line shows odor pulse duration. Color labels shown on the right.

**Extended data Fig. 4.**
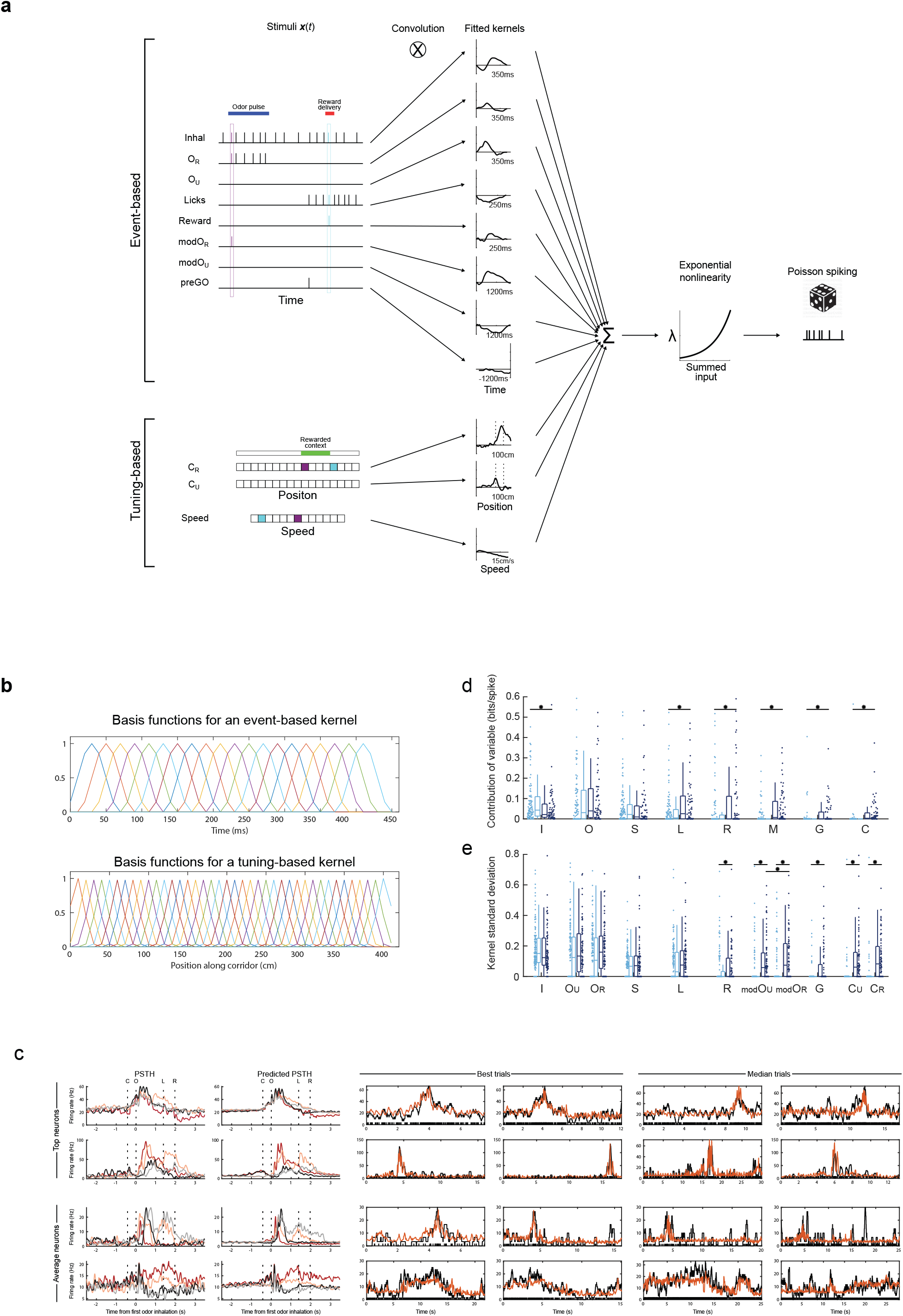
Parametrizations in GLM model. **a**, Scheme of parametrizations of task variables. Top: time evolution of variables parametrized for convolution with event-based kernels. Bottom: animal-state variables parametrized for convolution with tuning-based kernels. The instantaneous variable values in two individual 10-ms time bins are labelled in purple and cyan. **b,** Scheme of basic functions used to parametrize kernels: an example for an event-based kernel (top panel; inhalation kernel) and an example for a tuning-based kernel (bottom panel; spatial context kernel). **c**, Model-based PSTH and single-trial predictions. Examples of four neurons are shown (two “top neurons” and two “average neurons”, according to PSTH prediction accuracy). Each row shows the neuron’s measured and predicted PSTHs (first two columns) along with the measured and predicted single trials (last four columns, two “best trials” and two “median trials” according to trial prediction accuracy). For single trials, black ticks at the bottom indicate the observed spike train and the black trace shows an estimate of spike rate obtained by smoothing the spike train with a 300-ms boxcar, while the orange trace shows the model-based spike rate prediction for that trial. **d,** Contribution of each task variable to the GLM encoding model (bits per spike) in first and expert sessions. Black dots indicate statistically significant differences between first or expert session recordings (Wilcoxon rank sum test. I, p-value = 2.3 x 10-2; L, p-value = 2.6 x 10-2; R, p-value = 3.8 x 10-2; M, p-value = 6.9 x 10-8; G, p-value = 1.6 x 10-4; C, p-value = 8.12 x 10-18) **e,** Standard deviation of kernels as a function of each variable in first and expert sessions. Asterisks indicate statistically significant differences. Wilcoxon rank sum test was used for first session vs. expert session comparisons (top asterisks: R, p-value = 4.5 x 10^-2^; ModOU, p-value = 2.7 x 10^-7^; ModOR, p-value = 6.1 x 10^-8^; G, p-value = 1.1 x 10^-4^; CU, p-value = 6.4 x 10^-18^; CR, p-value = 5.2 x 10^-18^) and Wilcoxon signed rank paired test was used for comparison across kernel categories of each neuron (bottom asterisks: modOU vs modOR,p-value = 2.6 x 10^-2^).

**Extended data Fig. 5.**
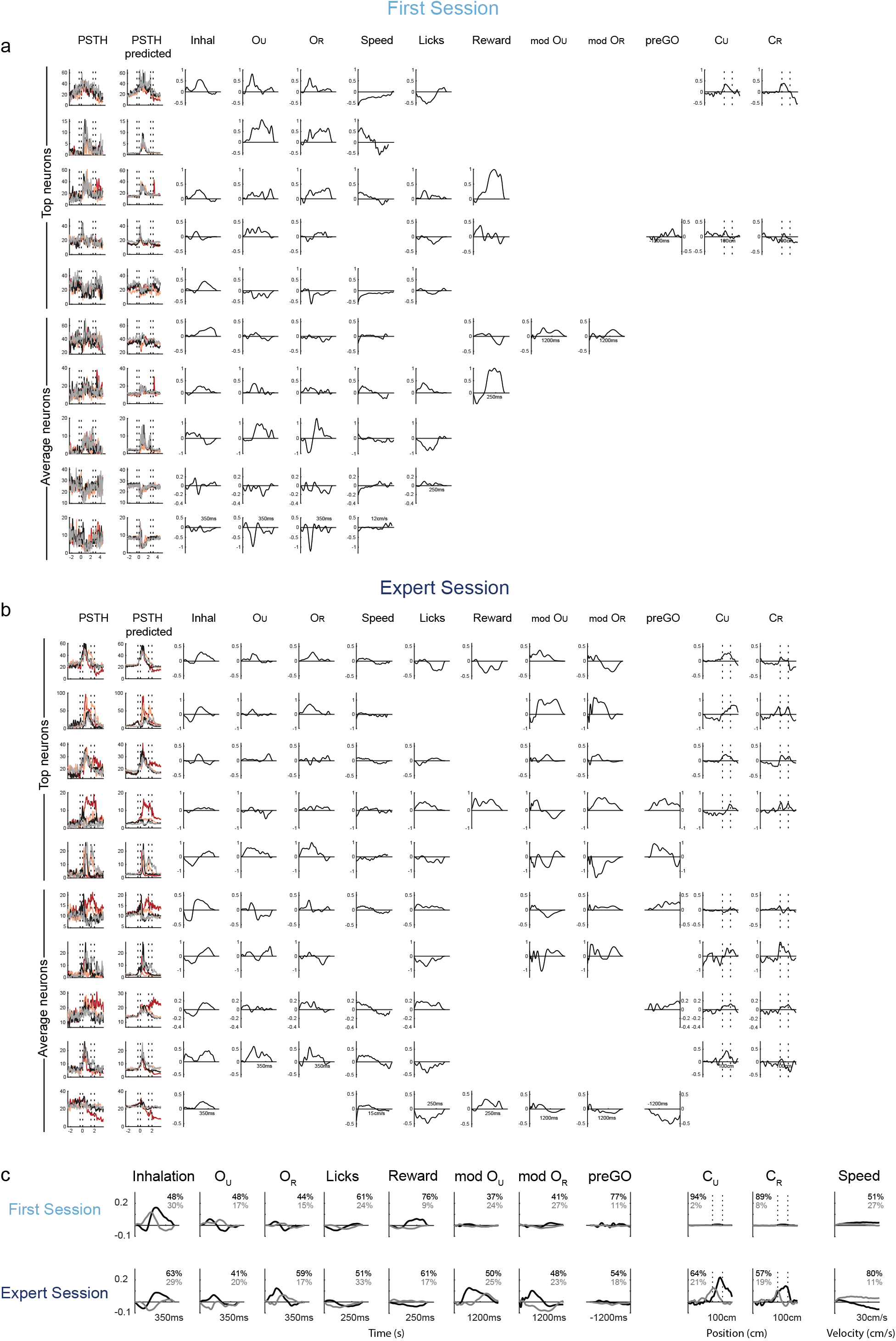
Examples of kernels obtained with the GLM encoding models of neurons. **a and b,** Examples of PSTHs and model-predicted PSTHs of several neurons recorded during first session (a) and during expert session (b). Panels on the right show the corresponding kernels obtained for each neuron. **c**, First (black) and second (grey) PCA component of pooled kernels for inhalation, unrewarded odour (OU), rewarded odour (OR), licking, reward, modulation of context (mod) onto OU, mod OR, activity before a GO decision (preGO), unrewarded context (CU), rewarded context (CR) and animal speed. Explained variance by each component is indicated in percentages, first-session recordings (top) and expert-session recordings (bottom). Parametrizations in GLM model.

**Extended data Fig. 6.**
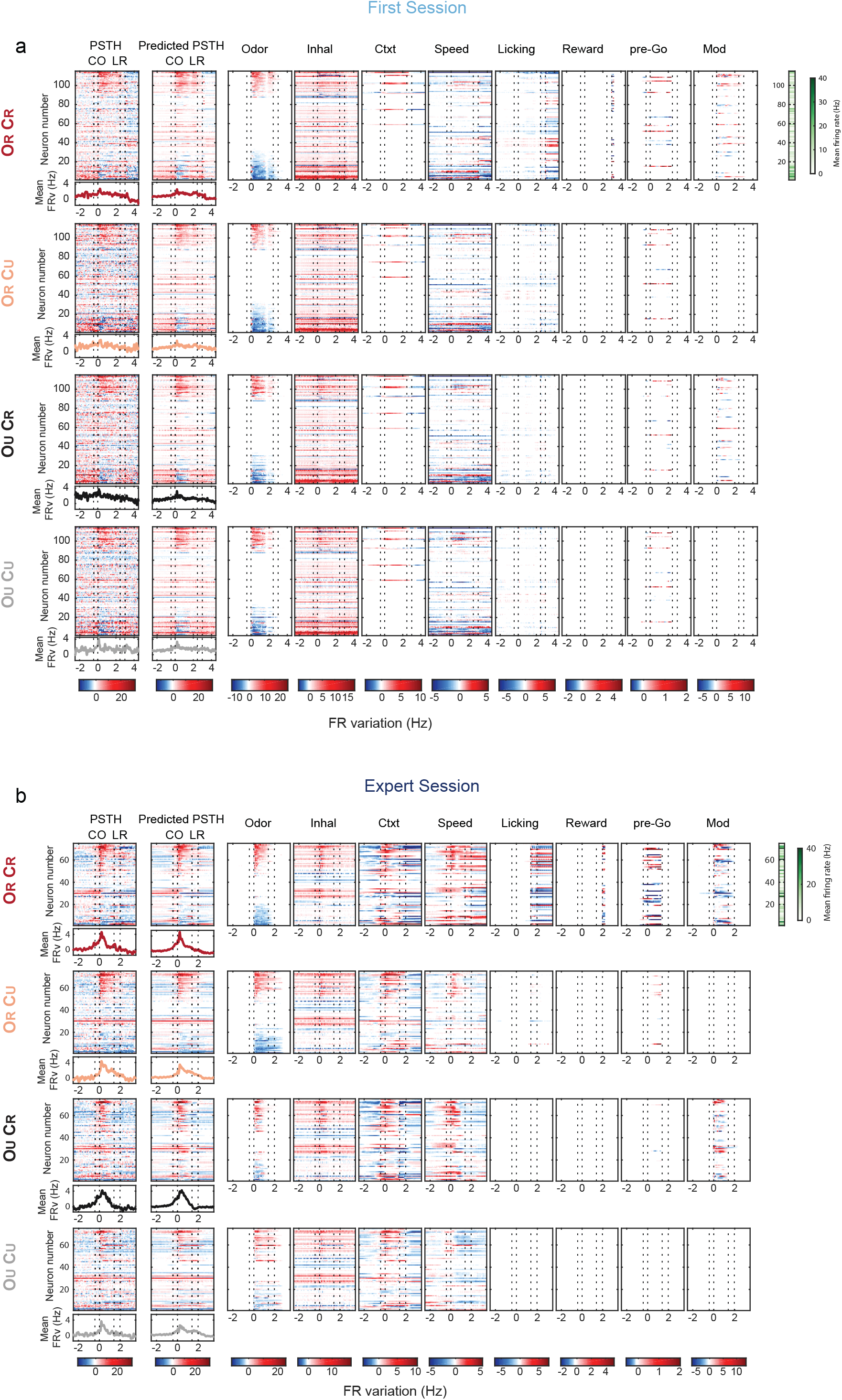
Firing-rate modulations induced by individual model variables. **a,** PSTHs and model-predicted PSTHs of all neurons with fitted kernels, recorded in first sessions. Panels on the right show predicted firing rate modulations (around the neuron’s mean firing rate) induced by each variable (i.e., for GLM models including only that task variable). Neurons were sorted according to their average responses to ORCR trials. **b,** Same as a but for expert sessions.

**Extended data Fig. 7.**
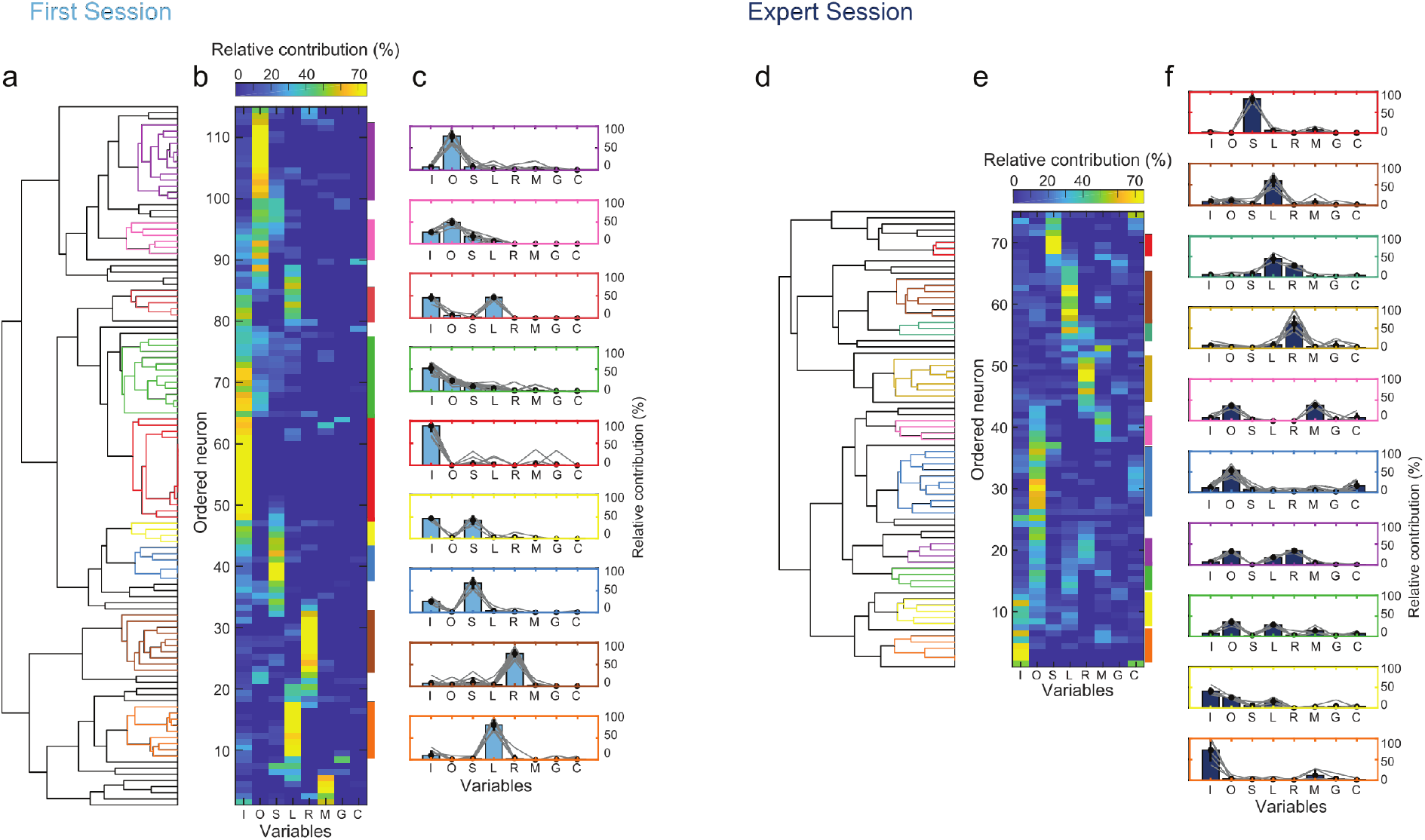
Clustering neurons according to the relative contribution of their encoded task variables. **a and d,** Dendrograms for the hierarchical tree of first-session (a) and expert-session (d) neurons sorted according to the relative contribution of their encoded task variables. **b and e,** Relative contributions of task variables to first-session (b) and expert-session (e) neurons, sorted according to the hierarchical tree. Colour bars on the right indicate each individual cluster obtained (same colour code applies to dendrograms). **c and f,** Bar plot for each individual first-session (c) or expert-session (f) cluster, showing the relative contribution of the variables encoded by the neurons in the cluster (same data shown in Fig. 3G).

**Extended data Fig. 8.**
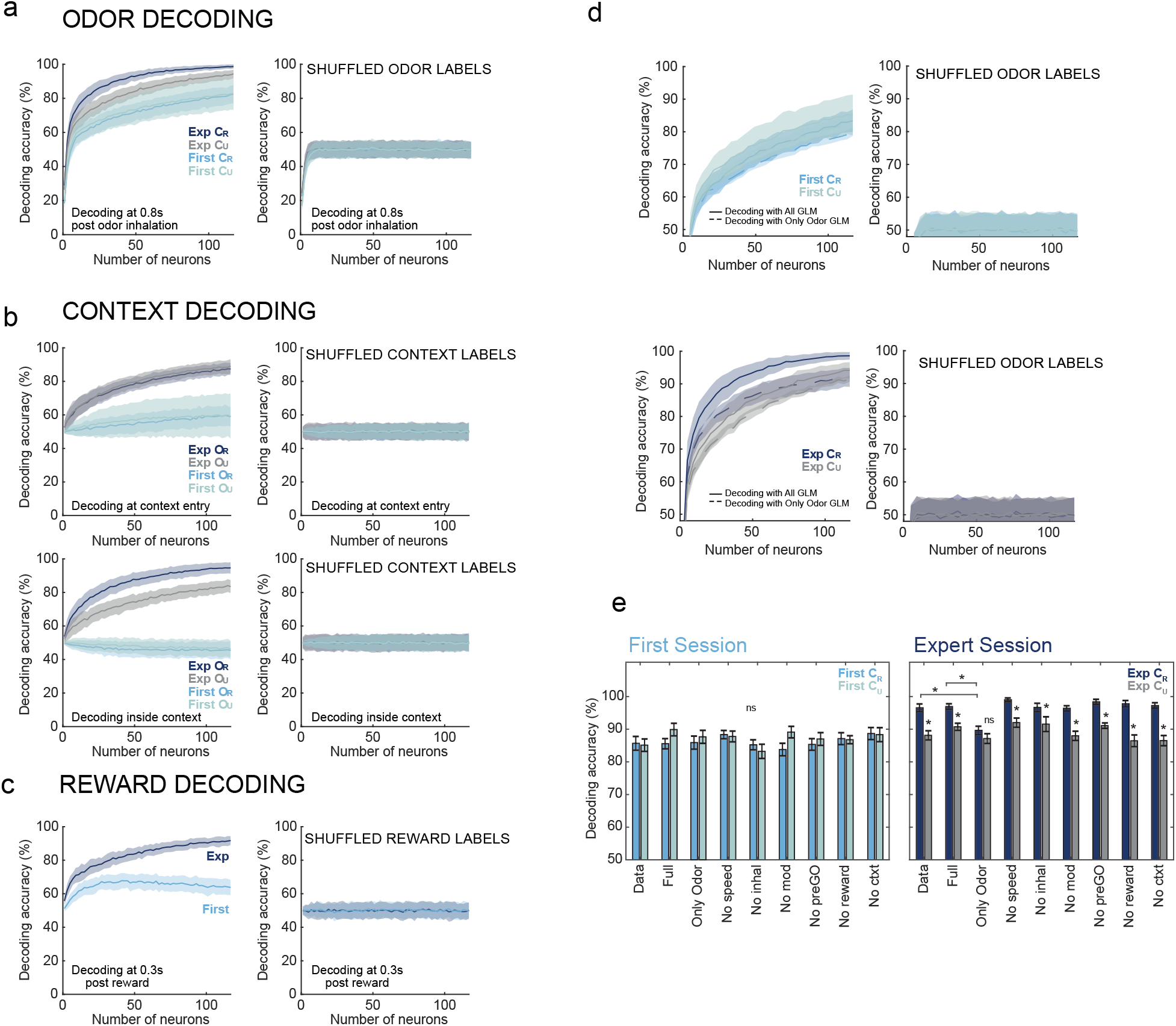
Multidimensional decoding in piriform neurons. **a-c,** GLM-based trial-by-trial decoding as a function of neuronal population size, using first-session (First) and expert-session (Exp) recordings, and after shuffling trial labels. **A,** Odour identity decoding at 0.8s after first odorant inhalation in rewarded and unrewarded context trials (CR and CU). **b,** Context identity decoding in rewarded and unrewarded odour trials (OR and OU). Top, decoding at the moment of context entry. Bottom, decoding when mouse is inside context zone. **c,** Decoding reward consumption 0.3s after reward delivery. **d**, Same GLM-based decoding analysis as in a, but decoding recorded data using GLM models including all fitted kernels (All GLM) and after removing contributions of non-odour-related kernels from the simulations of odour-responsive neurons (Only Odour GLM). Top, decoding accuracy for first-session data in CR and CU trials. Bottom, same as Top but for expert-session. Right panels show results after shuffling odour labels. For all decoding analysis shown in the figure the total number of trials used for training and testing GLM-based decoders was matched across the different trial types compared. Figure **e,** Decoding accuracy of linear classifiers for odour identity from data (Data) and GLM model simulations, in rewarded and unrewarded context trials (CR and CU, respectively), for first-session (First) and expert (Exp) animals. Odor identity was decoded with the activity of a population of 100 neurons, during a 0.5s window after first odorant inhalation. Simulations were obtained for GLM models including all fitted kernels (All GLM), GLM models where we removed from odour-responsive neurons all the non-olfactory kernels (Only Odour), and GLM models where we removed from odour-responsive neurons the following single kernels: context (No Ctxt), speed (No Speed), inhalation (No Inhal), modulation of odour responses by rewarded context (No Mod), anticipation of GO response (no PreGO) and reward consumption (No Reward). 2-Way Anova, for context trial type (CR and CU; 1 degree of freedom), Data type (Data and simulations from the 8 GLM model types; 8 degrees of freedom). Context-Data type interaction Prob>F: 2x10-05. One-way ANOVA corrected for multiple comparisons: Data CR vs Full CR, n.s.; Data CU vs Full CU, n.s.; Full CR vs Only Odor CR, p-value=2x10-06; Full CU vs Only Odor CU, n.s.; Full CR vs every GLM simulation with single kernels removed CR, n.s.; Full CU vs every GLM simulation with single kernels removed CU, n.s.; Data CR vs Data CU, p- value=7×10^-07^; Full CR vs Full CU, p-value=1×10^-04^; No Ctxt CR vs No Ctxt CU, p-value=7×10^-07^; No Speed CR vs No Speed CU, p-value=7×10^-06^; No Inhal CR vs No Inhal CU, p-value=7×10^-03^; No Mod CR vs No Mod CU, p-value=7×10^-07^; No PreGO CR vs No PreGO CU, p-value=2×10^-06^; No Reward CR vs No Reward CU, p-value=7×10^-07^; Only Odor CR vs Only Odour CU, n.s.

**Extended data Fig. 9.**
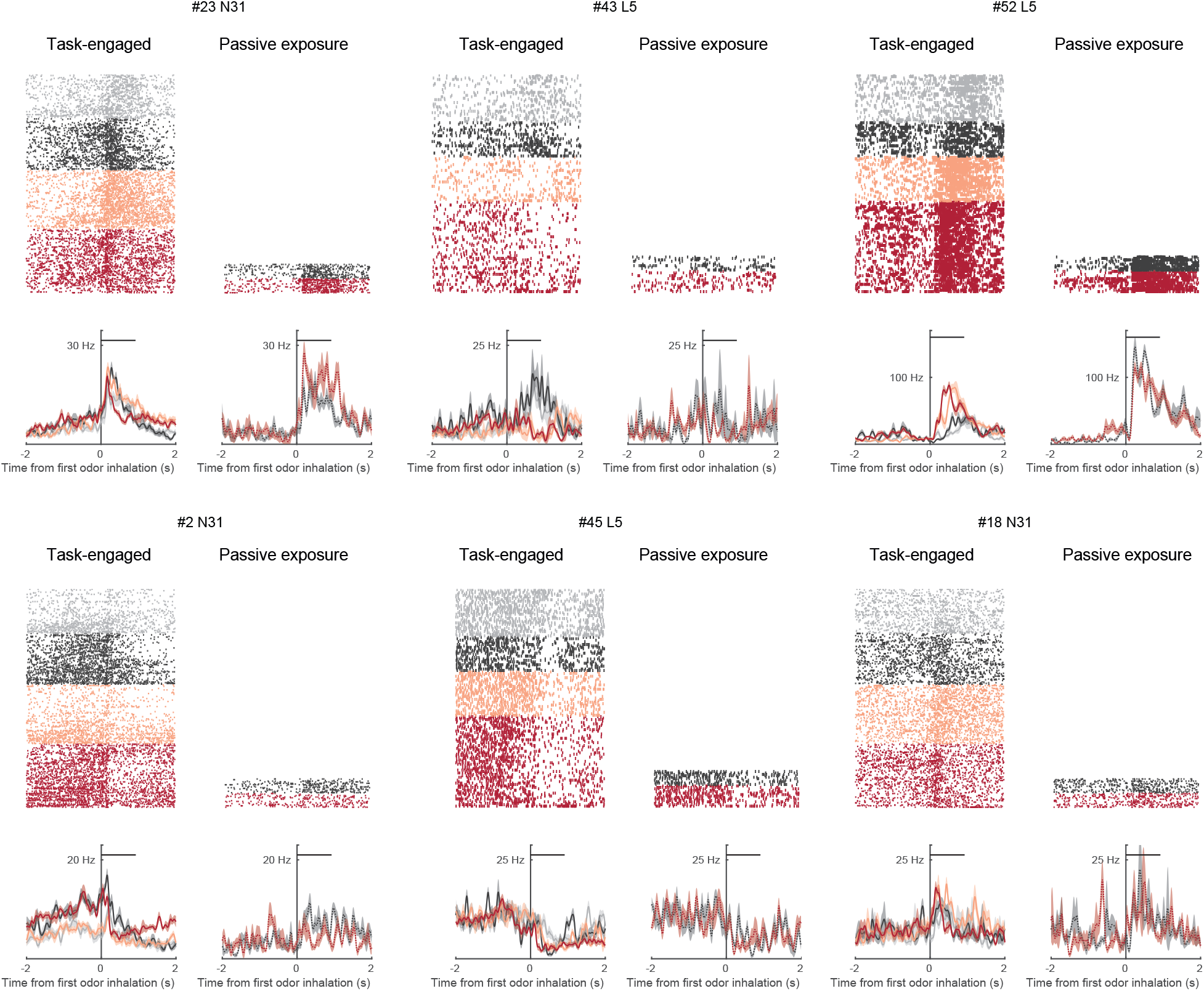
Passive vs task-engaged PCx responses. Example of neuronal recordings during task engagement or when the virtual reality was turn off and animals were passively exposed to odours.

**Extended data Fig. 10.**
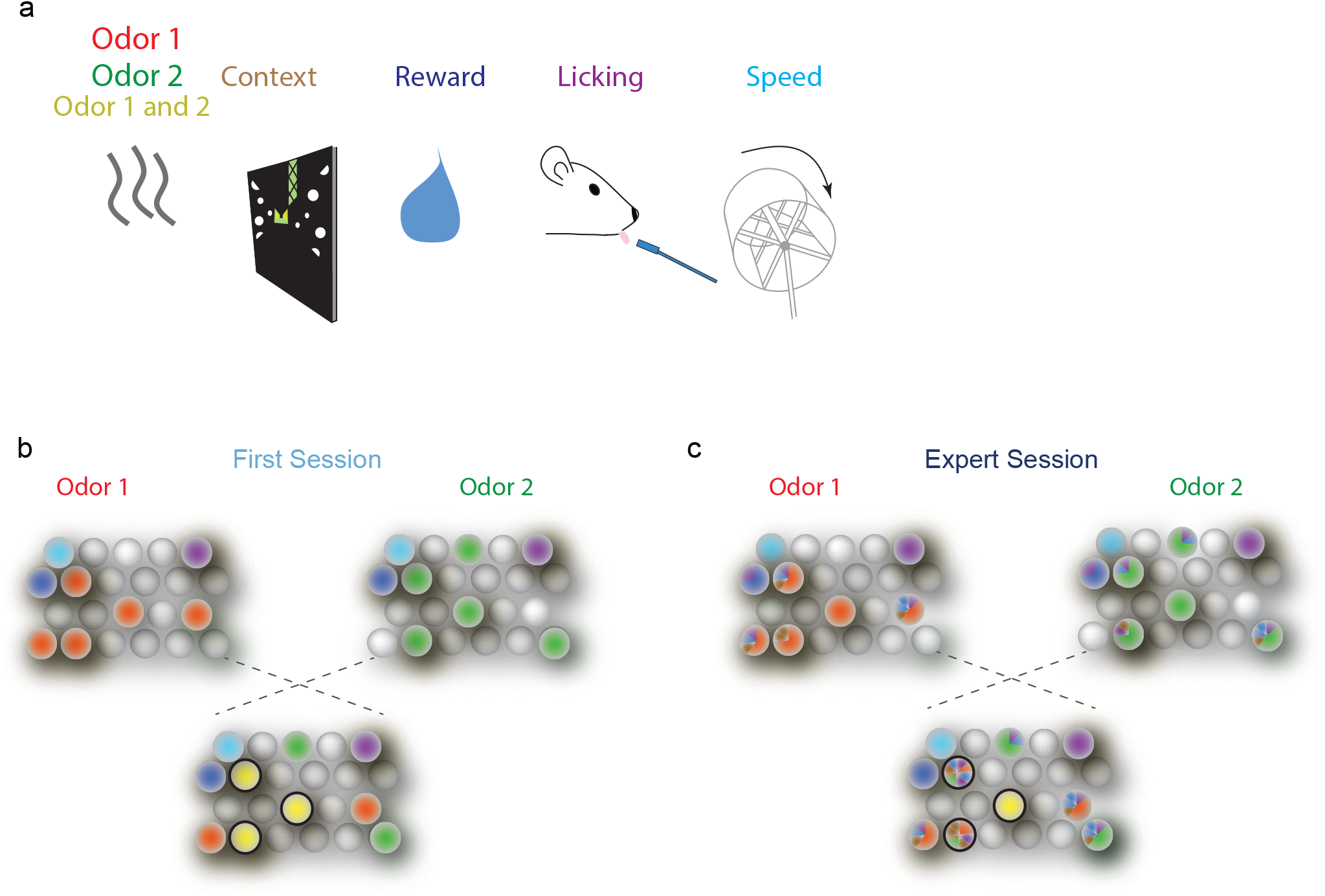
Learning induces mixed-selectivity in PCx neurons that improves discrimination. **a**, Variables associated with the odour-visual context-reward associative task. The variables are colour coded. **b,** The scheme represents responses of neurons to different variables from A, in an animal in the first session of training. Neurons tend to respond to single variables and do not show contextual modulation. Responses to individual odours have a percentage of overlapping illustrated by the yellow neurons that do not discriminate. **c,** The scheme represents responses of neurons to different variables from a, in an animal in the expert session of training. Neurons tend to have associative responses, with a contextual modulation. Notice that the overlap of the responses to the two odours decreases due to the mixed-selectivity that arises with learning the task.

## Methods

### Animals

#### Animals

We used 7-9 weeks old (n = 10 mice) female and male C57BL/6J mice. Six animals were pooled in the first-session group and four in the expert-session group. Blinding is not relevant to this study since all behavioral and neuronal activity analyses were performed with automated scripts without experimenter intervention or selection, and curation of spike-sorting results was done without knowledge of responsiveness to stimuli of each single-unit cluster. Littermates were randomly assigned to first-session or expert-session groups. No statistical methods were used to predetermine sample size. Mice were group-housed before head bar implantation and housed singly thereafter. The experimental timeline is shown in Extended data Fig. 1a.

#### Water Restriction Protocol

Water restriction started after mice recovered from surgery (at least three days post-surgery, Extended data Fig. 1a). Mice were individually housed in cages containing play tunnels, a wheel, and nesting material, in a reverse light cycle (12O:12L) housing room. The light in the accommodation room had an intensity of 100 lux at the height of the boxes, with white LEDs. The habituation, training and neuronal activity recording sessions were carried out during the dark phase. In the experimental device where the animals were placed, there was no illumination besides the light from the virtual reality screen. Relative humidity was maintained at 40-50%. Dry food was continuously available. One ml of water was dispensed manually into small containers attached to the inner walls of the cages, always at the same time of day. Mice were monitored daily for hydration level, weight, skin health, and locomotion. To control these parameters, a daily record of: movement in the box, cleaning/grooming, posture, tension/relaxation of the skin of the neck, defecation in the housing box was used. Our protocol (IBioBA-CICUAL # 2020-03-NE) was subject to rigorous animal welfare control measures, including daily weight determinations and activity scores, posture, grooming, intake, and signs of dehydration. In the event of a weight loss below the minimum body weight (70% of the initial weight), the total daily volume of water they receive was increased until the stabilization of the body weight in water restriction.

### Behavioral task

Mice were head-fixed and trained to run on a running wheel (a plastic cylinder) (Fig. 1a and Extended data Fig. 1b). A water spout was positioned near the snout of the animal, and licks were detected with an infrared beam and sensor. Wheel rotations as the animal ran were translated into displacements through a linear virtual corridor, which was presented in a screen in front of the animal. Mice were trained in a behavioral task of olfactory discrimination dependent on visual context. Here, visual context refers to the virtual environment, which could be one of two types of corridors. Each corridor consisted of an approaching aisle (83 cm in length), a context zone (33 cm in length) and a reward zone. Both types of corridors were visually distinct only in the context zone. The approaching aisle and the reward zone were similar for both corridors, with black walls displaying white spots. The context zone could be either green (green floor and walls with vertical white and green stripes, and a green column on the right with a black diamond-shaped pattern) or gray (black floor with white diamond-shaped pattern and gray walls with black dots, and a column on the left with horizontal white, black and gray stripes) (Extended data Fig. 1b). When the animals entered the context zone, they received an odor puff of 1-second duration through a pipe directed to the animal nose. The odorant could be of two types: isoamyl acetate or ethyl butyrate. Once animals left the context zone where they received the odorant stimulus, they entered the reward zone, where they could choose to lick (GO response) or not (NO-GO response) to obtain a drop of water reward. There are four possible odor-context combinations, only one of these was rewarded after the GO response in the reward zone (Fig. 1b). No punishment was given for incorrect GO responses or for licking elsewhere in the corridor in any trial type. After reward delivery or an incorrect GO response the trial was immediately finished and a new trial started by “teletransporting” mice to the start of the approaching aisle.

A specific rewarded odor-context combination (e.g., isoamyl acetate-gray context) was randomly assigned to each mouse and maintained throughout all training sessions. This rewarded combination was labeled O_R_C_R_. The individual odorant and context used in that rewarded combination were labeled O_R_ and C_R_ respectively, while the remaining odorant and context were labeled O_U_ and C_U_. The “R” and “U” subscripts stand for “rewarded” and “unrewarded”, but notice this is an abuse of language since specific odor-context combinations were rewarded, not individual odors or contexts (that is, only O_R_C_R_ trials were rewarded while O_R_C_U_ and O_U_C_R_ trials were not).

#### Passive stimulation

for 2 of the expert animals, at the end of the behavioral task we additionally performed a passive odorant stimulation protocol. The protocol consisted of 20 trials of stimulation with each odorant (trials were randomly interleaved), while the virtual reality monitor was turned off (Fig. 4b). Odors pulses lasted 1 second, and were presented every 40 seconds.

#### Experimental apparatus

The device consisted of: I. Columns for holding the animal’s head (head-fixed rig). Animals are fixed to the rig by clamping a skull-implanted metal bar (headbar) to the rig’s columns. II. A running wheel for the animal with a rotary encoder attached to the wheel’s axle that records wheel rotations as the animal walks (see *Data acquisition*). The wheel is made of plastic and is covered with black EVA rubber so that the substrate is soft to the touch of the animal and avoids slippage; III. A circuit of valves connected to pipes (olfactometer) that conducts the odor to the animal’s nose (see *Olfactometer* in *Odor Stimuli)*. This system is computer-controlled, which allows different odors to be presented with adjustable flow rates and to switch between odors quickly (<20 ms). In addition, an air exhaust quickly removes the odor after its presentation. Both the olfactometer and the exhaust allow the olfactory stimulus to be presented in a timely manner. Airflow is kept constant and regulated by flowmeters; IV. An inhalation-exhalation cycle measurement system: sniffing behavior was recorded with a mass airflow sensor located externally in close proximity to the animal’s left nostril (see *Data Acquisition* and *Analysis of respiration recordings*). Precise orientation relative to the nostril was manually optimized before each recording to attempt to acquire a full signal, despite any movement of the nose. V. A custom-made lickometer for measuring the animal’s licking response and for delivery of water rewards (see *Data Acquisition*). VI. A computer screen placed in front of the animal that displayed the virtual reality environment according to the task protocol. VII. A microcontroller and data acquisition system (Bpod State Machine r1, Sanworks) that records the animal’s licking response and its behavioral performance across trials, while controlling in real time the state of the task (activation of odor valves, delivery of reward, sequence of trial types) according to the task protocol. VII. A data acquisition system (Smartbox, Neuronexus) that acquires and stores behavioral and trial information, along with the electrophysiological signals (see *Data Acquisition*). IX. A CPU running MATLAB (Mathworks) for controlling the BPod and running the Virtual Reality MATLAB Engine, ViRMEn ^40^.

### Animal training

#### Behavioral training

After body weight stabilization, usually after five days of water restriction, the experimental device habituation and training session began (Extended data Fig. 1a). During those days, we carried out a daily *handling* or manipulation session in which the animals got used to the manipulation of the experimenter and to remain comfortable in the hands of the experimenter. *Habituation to the experimental device:* Three days before training, the mice were manipulated daily so that they got used to the environment of the training room and head fixation to the experimental device.

#### Behavioral task training

A random sequence of trial types (different odor-context combinations) was presented to the animal in each training session. Individual training sessions lasted 40-60 minutes. In the first training session animals completed 134 +/- 51 trials, while they performed 227 +/- 60 trials on expert sessions. Rewards were 10 microliter water drops.

### Odor stimuli

#### Olfactometer

Odors were delivered using a custom made olfactometer. Charcoal filtered air was routed into two flowmeters (03216-06 and 03216-16, Cole-Parmer, IL). The first one carried a neutral air stream at 0.9 liters per minute (LPM) that was kept constant (base stream). The second one carried a 0.2 LPM stream which was further splitted in two 0.1 LPM arms with an injection valve (SI360T041, NResearch, NJ; “odor bank valve”). When the valve was turned off the first arm was selected and the air stream traversed through a 10 mL empty vial containing (blank stream). When the valve was turned on the second arm was selected and the air stream was channeled to a second injection valve (“odor selection valve”) that routed the air to one of two 10 mL vials, each one containing a 2 mL solution of one of the two odors. The voltage state of this odor selection valve thus controlled which odor would compose an odorant stream. The two arms were combined in a shuttle valve (SH360T041, NResearch, NJ; “isolation valve”) that was synchronously activated with the odor selection valve to isolate both odor paths, avoiding odorant cross-contamination that could result from air reflux into the remaining vial. The odorant stream and the blank stream were then fed to a final shuttle valve that selected between both streams (“injection valve”). The selected 0.1 LPM stream (odorant or blank) was combined with the 0.9 LPM base stream to produce a 1 LPM stream that was routed to the animal nose. The Bpod system was programmed to control valve voltages so that when entering the visual context, the appropriate odorant stream would be selected and the animal would receive the 1-second-long odor pulse scheduled for that trial. Final valve switching simultaneously rerouted the odorant stream to the animal’s nose and the blank stream to a vacuum exhaust, and switched back after 1 second. At any other trial moment, the blank stream would be selected and the animal would receive an air stream that passed through vials containing only mineral oil. Non-selected odorant streams were also directed to the vacuum exhaust. Consistency of shape and arrival times of odor pulses were monitored with a custom-made photoionization detector (PID; PID-A1, Alphasense, Essex, UK) located at the outlet of the stimulation tubing near the animal’s nose. Calibration of the olfactometer was routinely performed such that switching of the injection valve produced minimal perturbations of the total air stream to the animal’s nose. Tubings were made of Teflon to avoid accumulation of residual odors.

#### Odorant stimuli

isoamyl acetate or ethyl butyrate (SIGMA-Aldrich). The odors were chosen based on literature showing that these two odors had no innate response in mice^19^. The odors mentioned were prepared in the liquid phase at room temperature and the volatiles of the gaseous phase were supplied to the animals as a stimulus. For all experiments, solutions with 1:200 dilutions were used (odor: mineral oil) for odorant vials. The volatiles present in the gas phase of the vials were administered to the animals through the olfactometer. Odorant and blank vials in the olfactometer contained a liquid phase of 2000 µl of the respective solutions and a gas phase of 10 ml in which vapors were accumulated until reaching equilibrium with their respective vapor pressure. As a source of purified air, an aquarium pump was used whose air was passed through activated carbon and cotton filters. Airflow was constantly controlled by valves driven by electrical controllers. The opening and closing of the air flow passing through each vial was regulated. When a valve was opened, a volume of the gaseous portion of the vial was displaced, mixed with a continuous main flow of purified air, and finally emptied into a tube with an outlet located 1-2 cm from the mouse’s head.

### Surgeries

#### Stereotactic targeting and head bar attachment surgery

Animals were injected intraperitoneally with Ketamine (100mg/kg body weight) and Xylazine (10mg/kg body weight). From the establishment of total anesthesia, controlled through the loss of body reflexes, the operation lasted a maximum of 30 min. It was done on a heating pad to preserve the animal’s body heat. The eyes were protected from drying out by applying ophthalmic gels such as Vidisic (active ingredient: polyacrylic acid). The entire procedure was performed under aseptic conditions. Once anesthetized, animals were placed in a stereotaxic. A dose of bupivacaine 0.5% (50µL) was injected under the skin as a local anesthetic, and we waited for 5 minutes before continuing. An incision was made in the skin to expose the skull, disinfecting the area with pervinox using sterile swabs. The animal was prepared to place a metal bar that allowed us to attach its head to the experimental device and keep it fixed, and on the other hand to mark the skull in the position where the recording electrode would be inserted afterwards. The scalp was cut, exposing the skull and the region to be drilled. The skull was cleaned and dried with sterile cotton swabs for better adhesion of the glue. The 2.5 cm x 0.5 cm, 350 mg weight aluminum headbar was placed directly on the wet glue and then dental acrylic was added to cover the glue and cement the bar in the desired position. The metal bar then stuck rigidly to the skull, without going through it. The bar was attached to the apparatus during behavior. With the aid of a mouse brain atlas, the desired position relative to the bregma in the left hemisphere (for piriform cortex AP 3.2mm; ML 3mm) was reached using a stereotaxic and a point was marked. We also implanted the ground and the reference electrodes (silver wires) in the cerebellum. After the operation, animals recovered on the heating pad until awakening from anesthesia. We have determined that within two hours after the operation mice wake up, walk, drink, eat and do not show behavior indicative of pain sensation. However, preventively, the analgesic Tramadol was administered in a subcutaneous dose (5 mg/kg body weight), and in the drinking water. In addition, an anti-inflammatory, Ketoprofen, was administered by subcutaneous injection (5mg/kg body weight) one dose, for two days. Animals were allowed to recover from surgery for 3 days.

### Neural recordings

In different training sessions, depending on the experimental group to which the animal belongs (first session or expert animals), the recording session was held, which consisted of presenting a sequence of different trials while recording the activity of neurons in the piriform cortex. To do this, one day before this session, the craniotomy was performed where the recording electrode was going to be acutely inserted the following day. For this, animals were anesthetized with isoflurane (2% induction, 0.5–1% maintenance). A drill was used to gently file down the bone in the marked area during the previous surgery. Then, with the help of a needle, a small “cap” was lifted, exposing a small area of the brain which was kept moist carefully using swabs moistened with saline solution, avoiding touching the brain. Finally, the exposed portion was covered with a thin layer of cyanoacrylic glue (WPI). At the end, 100 µl of an anti-inflammatory, Ketoprofen, was administered by subcutaneous injection (5mg/kg body weight), and then animals were allowed to wake up and recover. The next day, the awake animal was placed in the experimental device, the glue/gel covering the craniotomy was removed with forceps, and an array of recording microelectrodes (Silicon probes, Neuronexus) attached to a micromanipulator was inserted. With the aid of the micromanipulator we reached the piriform cortex, descending slowly (1 micron/second) to the target area (DV 3.8-5.2mm, depending on the animal). Once we arrived, we waited 20 minutes to stabilize the recording position. The recording probe was previously painted with a dye (DiI D3911, Thermofisher). All recordings were performed using A1x32-Poly3-5mm- 25 s-177 silicon probes (177 μm2 site surface area, 3-column honeycomb site geometry with 18 μm lateral and 25 μm vertical site spacing, 36 μm center-to-center horizontal span, 275 μm center-to-center vertical span, 114 μm maximum shank width near the sites, 15 μm shank thickness) with an H32 connector (NeuroNexus Technologies). The animal performed the behavioral task while the neuronal activity was recorded. Once the recording session was complete, the animal was placed in its housing cage with water ad libitum. The next day, it was anesthetized intraperitoneally with Ketamine (100mg/kg body weight) and Xylazine (10mg/kg body weight). Once anesthetized, without reflexes and in an unconscious state, the animal was decapitated and the brain dissected. The animal’s brain was used for verification of the location of the recording probe (Fig. 3a). To do this, the brain was cut into 50-micron slices on the cryostat, the slices were stained in DAPI preparations to visualize cell nuclei under a fluorescence microscope, and the location of the probe tip was revealed by the fluorescence of the dye DiI.

### Data acquisition

Electrophysiological signals were acquired with a 32-site polytrode acute probe (A1 × 32-Poly3-5mm-25s-177, Neuronexus, MI) connected to an Acute Smartlink32 headstage (Neuronexus, MI). Unfiltered signals were digitized at 30 kHz at the headstage and recorded by a Smartbox multichannel data acquisition system (Neuronexus, MI). Experimental events and respiration signals were acquired at 30 kHz by analog and digital inputs of the Smartbox system. Respiration was monitored with a microbridge mass airflow sensor (Honeywell AWM3100V, NJ) positioned directly opposite the animal’s nose. Negative airflow corresponds to inhalation and negative changes in the voltage of the sensor output.

Behavior was automatically monitored by a behavioral measurement system Bpod (Sanworks, NY), which was programmed to control task flow according to the protocol (see *Behavioral Task*) and sent behavioral timestamps using BNC signals to the Smartbox system for synchronization with neuronal signals.

Animal displacement on the running wheel was measured with an optical rotary encoder (H5-360-IE-S, US Digital, WA) attached to the wheel axle. When the animal moved the encoder sent TTL pulses to the Smartbox (for posterior synchronization with neuronal signals), and to a microcontroller (MEGA 2560, Arduino, NY) programmed to calculate animal position. The microcontroller transmitted the positional information to the Virtual Reality MATLAB Engine (ViRMEn). It also and TTL pulses to the Bpod to signal context zone entry (to trigger odor stimulation) and exit (to signal entry to reward zone).

### Data Analysis and Statistics

All data analysis and statistical tests were performed with custom-written software using MATLAB (Mathworks). Statistical significance was assessed with the Wilcoxon rank sum test, unless otherwise noted.

### Analysis of task performance

Task performance of the animals was quantified using Signal Detection Theory, SDT ^41, 42^. Briefly, through analysis of the Hit rate and False-Alarm rate of a subject performing a discrimination task, SDT allows to disambiguate the contributions of the perceptual strength of the stimulus and the subject’s bias to respond.

The parameter called *d’* (also *d-prime* or sensitivity) measures the perceptual strength of the target stimulus, that is the discriminability between the subject’s perceptual representations associated to the presence or absence of the target stimulus:

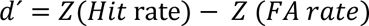

where *Z* is the normal inverse cumulative distribution function, and *Hit_rate_* and *FA_rate_* are the Hit and False-alarm rates, respectively. In our task, the target stimuli are the O_R_C_R_ trials. A value *d’* = 0 is chance performance, and higher values of *d’* indicate better performances. For example, for *Hit_rate_* = 95% and *FA_rate_* = 5% one obtains *d’* = 3.29 (dotted line in Fig. 1c).

The parameter called *criterion* (also *c*) is related to the subject’s bias response:

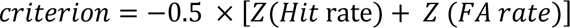

According to SDT, the subject decides to respond or not in a given trial by imposing a threshold: if the perceptual representation in the trial is larger than the threshold *criterion* then the subject responds with a “GO”. Negative criterion values indicate a bias towards *GO* responses, positive values indicate a bias towards *NO-GO* responses, and *criterion* = 0 indicates no bias (neutral response).

In our task, initial animal training sessions had negative *criterion* and low *d’* values, indicating almost chance performance and a bias towards licking across all trials (since animals were thirsty and there was no punishment for False Alarms). As training progressed, animals steadily increased *d’* (increasing performance) and brought *criterion* towards a neutral unbiased response (*criterion* → 0).

### Processing of respiration recordings

We took an approach similar to previous studies^43^. Respiration traces sampled at 30 kHz along with the neural recording signals were smoothed with a second-order Savitzky-Golay filter in 100 ms frames and the result was locally detrended by subtracting its 1-second-long median-filtered signal. The start of inhalation was defined as zero-crossings before large negative peaks in the smoothed, detrended signal. Odor inhalations were defined as inhalations happening during a 1-second long time window starting immediately after the arrival of the odorant pulse.

### Spike Sorting

Extracellular voltage traces were preprocessed with common median referencing (subtraction of the median across all channels at each time sample to remove artifacts), spike sorted using Kilosort^44^ (https://github.com/cortex-lab/Kilosort) and the obtained result was manually curated with the phy GUI (https://github.com/kwikteam/phy). During manual curation all clusters of putative spike events detected by each template were inspected and evaluated according to a number of criteria. First, we discarded clusters that were considered noise if their events had near-zero amplitude or if event waveforms were non-physiological and/or extended across all recording channels. Clusters containing inconsistent waveform shapes or large number of refractory period violations (<2 ms) in the autocorrelogram were also discarded. In a final step we merged pairs of clusters that had similar waveforms, showed refractory period cross-synchronization or temporally coordinated cross-correlograms that indicated a bursting neuron. Units that passed these criteria were labeled as single units and considered in our study.

### Single-neuron responses

For plotting and analysis of neuronal responses we used peri-stimulus time histograms (PSTHs) that were temporally smoothed with a gaussian filter (s.d., 30 ms). These “event PSTHs” described the neuronal spiking rate temporal evolution around the timing of either the onset of sensory stimuli, task events or behavioral events (examples in Fig. 3 and Extended data Fig. 3). For describing average responses along a complete trial we constructed PSTHs that were obtained by stitching together the “event PSTHs” corresponding to the following sequence of trial events: visual context entry, first odor inhalation, animal’s response, and trial outcome (labeled as “C”, “O”, “L” and “R” in Figs 1e-f, 2c and 3d). The “animal’s response” trial event refers either to the timing of the first animal lick after odor delivery (in GO trials) or to the timing at which the animal leaves the visual context (in NO-GO trials). The “trial outcome” trial event refers to the timing of reward delivery (in HIT trials) or to the timing that corresponded to an additional median inter-lick time interval after the “animal’s response” trial event (in MISS, FALSE ALARM and CORRECT REJECTION trials; the rationale for this is that rewards were delivered in the second lick performed in the reward zone). Each one of these 4 “event PSTHs” (C, O, L and R) were aligned to each event median time across trials, relative to first odor inhalation (for first session data C, O, L and R were aligned to -0.46s, 0s, 2.45s and 2.99s, respectively; for expert session data C, O, L and R were aligned to -0.43s, 0s, 1.38s and 1.96s, respectively). The final PSTH that described average activity of a neuron along a full trial was obtained by stitching together this sequence of time-aligned “event PSTHs” by averaging periods where there was temporal overlap between them. We confirmed that this stitching procedure did not distort the shape of the individual “event PSTHs” and precisely described the sequence of firing rate variations along the trial. All stitched PSTHs are shown in Fig. 1e, f and Extended data Fig. 6 and (where the mean firing rate of each neuron was subtracted to display variations in firing rate around the mean).

### Principal Component Analysis (PCA) and Demixed PCA (dPCA)

Dimensionality reduction of the population activity data was performed with PCA or dPCA^25^. The dPCA analysis was performed with the Matlab implementation provided by the authors (http://github.com/machenslab/dPCA). Details on dPCA can be found in the original publication, here we give a brief outline and describe how we applied dPCA to our data.

The data was organized by pooling all the full-trial stitched instantaneous firing rates across recordings and stimulus conditions, resulting in one array *X* for each training stage (i.e., first-session or expert-session stage). The dimensions of *X* were *N* x *C* x *O* x *T x K*, where *N* is the total number of neurons across recordings, *C*=2 is the number of contexts, *O*=2 is the number of odors, *T* is the number of firing rate time samples, and *K* is then number of trials. Thus, for each training stage, the array *X* has *N* rows in which the *i*-th row contains the instantaneous firing rate of the *i*- th neuron for all stimulus conditions and all trials (the firing rates were centered, i.e., with row means subtracted). By averaging *X* over all *K* trials for each neuron, odor, and context, we obtained a matrix 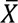 of dimensions *N* x *C* x *O* x *T* collecting the centered full-trial stitched neuronal PSTHs. For PCA, the 4-dimensional array 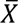 was reshaped into a matrix 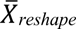 with *N* rows and *M* columns, where M= 𝐶 · 𝑂 · 𝑇 (that is, concatenating all stitched PSTHs conditioned on odor-context combinations), and singular value decomposition was applied on 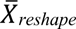 to obtain the matrix *W* of principal components. Trajectories through principal component space were obtained performing the projection 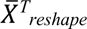 * W, where ^T^ refers to the transpose, reshaping this projection back to the 4-dimensional *N* x *C* x *O* x *T* form and plotting only the first 3 rows that correspond to trajectories through the top 3 principal components (Fig. 2b).

The aim of dPCA is to reduce the dimensionality of the data, while obtaining latent components that “demix” task parameters. For this, the single-trial data array *X* was reshaped into a *X_reshape_* with *N* rows and *P* columns, where P= 𝐶 · 𝑂 · 𝑇 · 𝐾 . When shaped this way, *X_reshape_* can be linearly decomposed into a set of 4 marginalizations 𝑥_Ф_: one condition-independent term, and 3 additional terms containing firing rate variations specifically related to either odorant identity, visual context identity, or the interaction between odorant and visual context identities. Each of these marginalizations 𝑥_Ф_can be understood as class means of the data, associated with the task parameter Ф. Importantly, these terms are all uncorrelated, thus the covariance matrix *X_reshape_X^T^_reshape_* can also be linearly decomposed into the sum of covariance matrices 𝑥_Ф_𝑥_Ф_^𝑇^corresponding to each of the 4 individual marginalizations. dPCA thus seeks to find separate encoding and decoding matrices that, when applied to *X_reshape_*, they minimize the least-squares reconstruction error of each individual 𝑥_Ф_simultaneously. It can be shown that this minimization is a reduced-rank regression problem that can be analytically solved by SVD (hence its relation to PCA). This dimensionality reduction step allows to infer a few latent components, also referred as decoding axes, that “demix” each task parameter and can be used as linear classifiers to properly decode the corresponding parameter throughout time in single trials (Fig. 2c and Extended data Fig. 2). Cross-validation was used to measure the time-dependent trial classification accuracy, and the significance of this accuracy was assessed by comparing performance against a shuffled set of trials (black horizontal bars in Fig. 2c and Extended data Fig. 2c, d indicate periods of significant classification accuracy). To avoid overfitting with dPCA we included the regularization term controlled by a parameter chosen through cross-validation on each dataset (results obtained without regularization were qualitatively similar).

In order to decompose data over a specific set of parameters, dPCA requires the data to contain trials for all possible parameter combinations and all neurons. Therefore, we could not further decompose our data over a decision (GO / NO-GO) parameter, since some parameter combinations were not available in the recordings (e.g., across expert-session recordings only one animal produced NO-GO responses in O_R_C_R_ trials and that same animal did not produce GO responses in O_U_C_U_ trials). Nevertheless, clear decision-related information could still be observed in our dPCA data decomposition, particularly in expert recordings, as departures of O_R_C_R_ trajectories from the rest of the conditions around the animal response time. This was possible since 90.3% of GO responses (243 out of 269) were concentrated in O_R_C_R_ trials when animals become experts.

### Generalized Linear Model (GLM) for encoding and decoding analysis

For this analysis the spiking-activity time series were discretized into 10-ms bins. The analysis was performed by adapting the Matlab implementation of Poisson GLM regression for trial-based spiking data by Park and colleagues^29^ (https://github.com/pillowlab/neuroGLM).

This framework poses an encoding model *p*(***r***|***x***) of the probabilistic relationship between a neuronal response **r** and a set of task variables ***x*** in a given trial. The probability of response is related to an underlying time-varying spiking rate λt, which is given by:

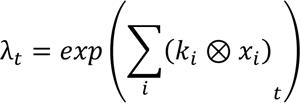

where 𝑥_𝑖_ (𝑡) is the time course of *i*th variable in the model, 𝑘_𝑖_ is the neuron’s “kernel” (i.e., linear filter) associated with that variable, and ⊗ indicates a linear convolution operation. The kernel captures the relationship between this variable and the neuron’s probability of spiking. In turn, in a given single trial, the probability of spiking of the neuron is given by a Poisson distribution (i.e., it assumes Poissonian spiking statistics):

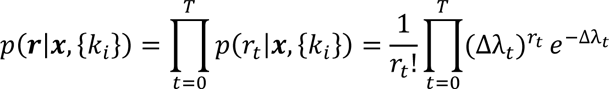

where Δ is the time bin size (10 ms), *T* is the total number of time bins in the trial, and 𝑟_𝑡_ is the spike count at time bin (*t, t+*Δ).

#### Parametrization of model variables and kernels

A schematic representation of the parametrization is shown in Extended data Fig. 4a. This choice of regression variables was based on the neuronal response modulations observed in the PSTHs and in population dynamics (dPCA). In our model there were two kinds of kernels: “event-based” kernels (i.e., kernels reflecting the time-varying influence of a task variable on the time-varying spike rate), and “tuning-based” kernels (i.e., kernels describing the dependence of neuronal firing on the value of a task variable). Kernels were parametrized by a set of basic functions (Extended data Fig. 4b). For any kernel parametrized by *n* bases, the model infers a set of *n* coefficients that correspond to the weighted linear combination of the *n* basis functions. To choose the number of bases *n* used to parametrize each kernel, we ran a series of preliminary tests on a set of neurons and evaluated parametrization performance through cross-validation (see *Model fitting and model performance*).

##### i. Event-based kernels

Each event associated with the variable was represented as a delta function over time, and convolved with the corresponding kernel (Extended data Fig. 4a). To parametrize the associated event kernels, we used basis functions defined by a series of raised cosine bumps centered at different times spanning the range of time covered by the kernel (Extended data Fig. 4b).

● Inhalation onset: 10 bases covering 460 ms window after the onset of each inhalation. Labeled “Inhal”.
● Inhalations of rewarded and unrewarded odors: 10 bases covering 460 ms window after the onset of each odor inhalation during odor stimulation. One kernel for each odor, labeled “O_R_” and “O_U_”.
● Licking: 15 bases covering 360 ms window after the onset of each lick. Labeled “Licks”.
● Reward consumption: 15 bases covering 360 ms window after the onset of each lick. Labeled “Reward”.
● Anticipation to GO response: 22 bases covering 2000 ms before the first lick that follows odorant stimulation. Labeled “preGO”.
● Modulation of odor responses by presence of rewarded context: 20 bases covering 1500ms after the first odor inhalation. One kernel for each odor, labeled “modO_R_” and “modO_U_”. These kernels aim at describing the interaction between odors and contexts.

##### ii. Tuning-based kernels

Each variable was discretized and represented as an animal-state vector for the variable at a given instant in time, and convolved with the corresponding kernel (Extended data Fig. 4a). Each animal-state vector denotes a binned variable value, all of whose elements are set to 0, except for one element, which is set to 1, corresponding to the bin the current animal state occupies. To parametrize these tuning kernels, we used basis functions defined by a series of raised cosine bumps centered at different variable values along the range spanned by the animal-state vector (Extended data Fig. 4b).

● Animal position along rewarded and unrewarded visual contexts: 42 bases covering the entire length of the virtual corridor discretized in a sequence of 4-cm bins. One kernel for each context, labeled “C_R_” and “C_U_”.
● Running speed: a sequence of bins of 1 cm/s up to the animal’s maximum running speed (the total number of bases varied between animals depending on their maximum speed, typically ∼35 bases were used). Labeled “Speed”.

##### iii. Bias kernel

To account for the average firing rate of the neuron during the experiment, we introduced a bias kernel parametrized by a single coefficient that was convolved with a boxcar function that lasted for the whole recording (that is, the bias kernel was constantly active). This kernel is not shown on Extended data Fig. 4a.

#### Model fitting and model performance

To fit the weights of the basis functions that parametrize the kernels in the GLM encoding model of each neuron, we maximized the model’s log-posterior, using a ridge prior to regularize the inferred weights (under the assumption that kernels should be relatively smooth):

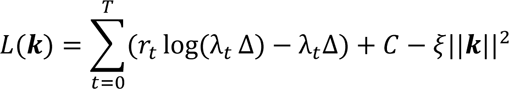

where ***k*** is a vector representing the weights to be obtained, *C* is a constant that does not depend on ***k*** (thus irrelevant for the optimization procedure), and 𝜉 is the smoothing hyperparameter controlling regularization.

Model performance was quantified through cross-validation, by computing the model log-likelihood (the log-posterior without the last regularizing prior term) of held out data under the model. For this, we randomly divided trials into 10 sets of equal size. The cross-validation procedure was repeated on each of these 10 sets of trials (10-fold cross-validation; the log-likelihood of the model on each fold was normalized by the number of spikes in the fold). This procedure penalized model overfitting, thus allowing valid performance comparisons between models of varying complexity.

To select the hyperparameter 𝜉 we took an Empirical Bayes approach, in which we maximized the marginal likelihood (also called model evidence) with respect to 𝜉 (chosen from a grid) and used this estimate of 𝜉 into the posterior. To compute the marginal likelihood we used the Laplace approximation. This procedure does not require cross-validation, and it was performed on all trials at once.

Once a model was selected (see *Model selection*), the final estimate of ***k*** was inferred by fitting the model on all trials at once.

#### Model selection

Three variables were associated with 2 kernels that depended on the identities of odorants and/or contexts: inhalations of rewarded and unrewarded odors (O_R_ and O_U_), modulation of odor responses by presence of rewarded context (modO_R_ and modO_U_), and position along rewarded and unrewarded visual contexts (C_R_ and C_U_). When regressing neuronal responses against these 3 variables, the 2 corresponding kernels were used. Thus, the total number of regression variables was 8, and the models considered included combinations between all of them.

Testing all combinations of variables is intractable. Thus, we implemented a forward-search model selection procedure similar to the one introduced by Hardcastle and collaborators^45^, that selects the simplest model that best predicts held out neuronal activity data (spike-normalized model log-likelihood on 10 cross-validation data folds; see *Model fitting and model performance*). First, models including a single variable were evaluated, and the one with highest performance was determined. This single-variable model was then compared against all models with two variables that included this single variable. If the double-variable model with highest performance was significantly better than the best single-variable model, we proceeded to compare this double-variable model to the best performing triple-variable model that included this pair of variables. We continued including variables in the model through this procedure until the more complex model considered did not show an improvement in prediction performance, in which case the simpler model was selected. Significance in performance improvement was always assessed with a two-sample one-sided signed rank test, which tests the hypothesis that the difference in performance across data folds between the more complex and the simpler model comes from a distribution with median greater than 0. Models were considered better performing if they had a *p*-value lower than 0.05 under this test. Neurons for which the selected model did not perform significantly better than a model with constant mean firing rate were considered as not modulated by task variables (i.e, they had 0 modulating variables).

#### Contribution of a task variable to the GLM encoding model

We estimated the contribution of a given variable 𝑥_𝑖_ (associated with a kernel 𝑘_𝑖_) to the encoding model of a neuron by calculating the difference in model performance (spike-normalized log-likelihood increase) between the selected model θ (with firing rate λ_𝑡_) and the model 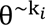 (with firing rate 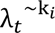) that contains all variables in the selected model, minus the variable 𝑥_𝑖_:

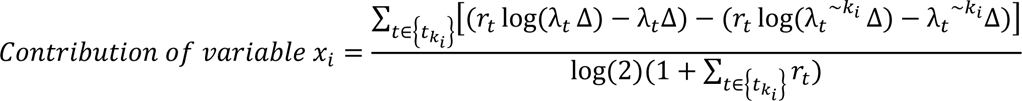

Here the sum is performed over the set of time bins {𝑡_𝑘_𝑖} in which the convolution (𝑘_𝑖_ ⊗ 𝑥_𝑖_) was performed, that is the moments in time were the kernel 𝑘_𝑖_contributed to the firing rate λ_𝑡_. The normalizing factor has an offset of 1 in the sum of 𝑟_𝑡_ over {𝑡_𝑘_𝑖} to avoid numerical undefined expressions when no spikes were emitted during {𝑡_𝑘𝑖_ }, and the log(2) is included to measure the spike-normalized log-likelihood increase in terms of bits. The *Contribution of variables* calculated for all recorded neurons is shown in Extended data Fig. 4d. For the percentual *Relative contribution* of each variable (Fig. 3e), for each neuron we normalized each variable contribution by the sum of all contributions:

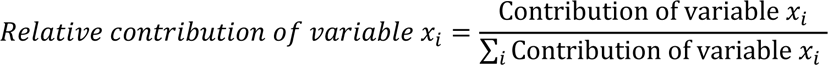

#### Decoding with GLM model

Decoding with the fitted GLM models, allows trial decoding of binary variable categories, while taking into account neuronal response modulations related to all the other task variables. As was previously shown^35^, under this GLM model, the computation for decoding the identity of a variable 𝑥_𝑗_ with two categories 𝑥_𝑗_ = 𝐴 and 𝑥_𝑗_ = 𝐵 (e.g., rewarded or unrewarded odor) with kernels 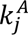 and 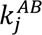, respectively, is rather straightforward. Consider the log likelihood ratio (*LLR*) for this situation:

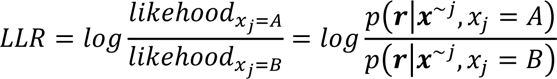

where 𝒙^∼𝑗^ is a vector with the rest of the variables other than 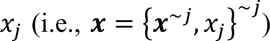. Further developing this formulation leads to:

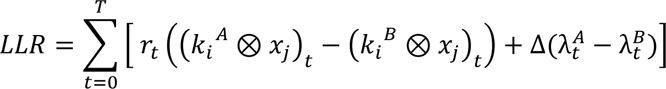

where 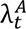 and 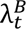 are the time-varying spiking rate for the model with 𝑥_𝑗_ = 𝐴 and 𝑥_𝑗_ = 𝐵 respectively. The *LLR* leads to a simple rule for decoding the category of the variable 𝑥_𝑗_: when *LLR* > 0, the recorded neuron indicates that 𝑥_𝑗_ = 𝐴 that is more probable under the model; on the contrary, 𝑥_𝑗_ = 𝐵 is more probable if LLR < 0. This mathematical procedure can be conceptualized as a time-varying estimation of whether the emitted spikes up to that time point in the trial are more probable under the spike rate prediction of a model that considers one of the variable categories rather than the alternative. When considering a population of neurons recorded simultaneously, the *LLR*s of the individual neurons can be summed together. In Fig 4a, this decoding accuracy is averaged across animals. For the analysis of the relation between the accuracy and the number of neurons used for decoding (Extended data Fig 8a-c), pseudopopulations of neurons were constructed with subsets of neurons that were randomly sampled from across recording sessions, up to the total number of 117 neurons (matching the total number of neurons from first-session and expert-session recordings). For each population size, the sampling procedure was repeated 50 times and accuracy was averaged. For the analysis shown in Extended data Fig. 8d aimed at evaluating the contribution to odor decoding accuracy of non-olfactory information encoded in PCx neurons, we calculated *LLR*s where *A* was set to *O_R_* and *B* to *O_U_*, and 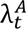 and 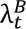 were calculated in two different ways: one using the full models (i.e., including all fitted kernels) and another after removing contributions of non-odor-related kernels from the models of odor-responsive neurons (i.e., removing non-olfactory modulations from neuron models that had olfactory kernels).

Expert-session recordings typically contained more trials than first-session recordings. To control for the effect of this difference on model estimation and hence accuracy of decoding, we re-fitted GLM models after matching the number of trials used from both training conditions. All decoding results are shown after this trial matching procedure.

### Decoding odors with linear classifiers

For the odor decoding analysis of Passive vs. Task conditions (Fig. 4c) and the evaluation of the contribution to odor decoding accuracy of non-olfactory information encoded in PCx neurons (Fig. 4d), we used linear classifiers based on L2-regularized logistic regression with 3-fold cross-validation. This analysis is independent of the GLM models or GLM-based decoding. This guarantees that the obtained decoding accuracies will not be influenced by differences across conditions in the goodness-of-fit or expressive power of GLM models.

Neuronal responses were concatenated into a Trials (T) x Neuron (N) matrix, arranged so that each row contained trial responses to a particular odor. Each neuron’s response was averaged over a 0.5-s window after first odor inhalation (for the Passive condition, peak decoding was reached later, at 1s, so we used that time window). Trials were labeled according to the odor sampled, and arranged into a vector of T elements.

Pseudopopulations of neurons were constructed with subsets of neurons that were randomly sampled from across recording sessions, up to the total number of 117 neurons (matching the total number of neurons from first-session and expert-session recordings). Because different trial conditions contained different numbers of trials, a random subset of responses from each neuron was excluded so that the number of trials per condition was consistent across neurons in the pseudopopulation. For each population size, this trial sampling procedure was repeated 50 times and the obtained accuracies were averaged.

For the analysis of Fig. 4d, a total number of 48 trials were used. For the Passive vs. Task conditions a total number of 25 trials were used (the fewer number of trials explains the lower performance reached in Fig. 4c compared to Fig. 4d).

### Clustering groups of neurons with similar relative contributions of their encoded variables

To group neurons according to the task variables that modulate their activities, we used an agglomerative hierarchical clustering approach. We only considered neurons whose GLM encoding model contained at least one task variable (i.e., with a total number of modulating variables >= 1; first session: 115 out of 177 neurons, 65.0%; expert session: 75 out of 117 neurons, 64.1%). First, for each neuron *k* we determined the vector 𝑹𝒆𝒍𝑪𝒐𝒏𝒕𝒓𝒊𝒃_𝑘_ = {𝑅𝑒𝑙𝑎𝑡𝑖𝑣𝑒 𝑐𝑜𝑛𝑡𝑟𝑖𝑏𝑢𝑡𝑖𝑜𝑛 𝑜𝑓 𝑣𝑎𝑟𝑖𝑎𝑏𝑙𝑒 𝑥_𝑖_}_𝑖_containing relative contributions of all variables to the neuron *k*. We then used a correlation metric to calculate the pairwise distances between all vectors of relative contributions, 𝑝𝑑𝑖𝑠𝑡_𝑖,𝑗_ = 1 − r_𝑖,𝑗_, where r_𝑖,𝑗_is the sample correlation between the pair (***RelContrib****_i_*, ***RelContrib****_j_*). With these distances we built an agglomerative hierarchical cluster tree, where distances between clusters were specified by the unweighted average distance between all pairs of objects in any two clusters (UPGMA). The obtained hierarchical trees are shown in Extended data Fig. 7a, d, and the vectors of relative contributions sorted according to their corresponding clusters are shown in Extended data Fig. 7b, e. To cut a hierarchical tree into individual clusters, we grouped tree leaves using a cutting threshold *thresh_cut_* for the distance coefficients of nodes in the tree (i.e., cutting through the tree at nodes whose heights were lower than *thresh_cut_*; clusters with less than 3 neurons were discarded). The obtained clusters of relative contributions are shown in Fig. 3g, Extended data Fig. 7c, f with a color code that is maintained in Extended data Fig 7.

We determined the optimal value for *thresh_cut_* by cross-validation of the generalizability of the resulting clusters. In order to do this, we randomly divided trials into 2 sets of equal size (sets *A* and *B*) and separately fitted the GLM model to each trial set, obtaining the vectors of the relative contributions of all variables to neuron *k* under the set *A* and under the set *B*, ***RelContrib^A^****_k_* and ***RelContrib^B^****_k_* correspondingly. We first evaluated how well clusters in set *A* generalized to set *B*: we labeled *A* as the train set, and *B* as the test set. We then fixed *thresh_cut_* on a given value, and performed the described hierarchical clustering procedure for both ***RelContrib^A^****_k_* and ***RelContrib^B^****_k_*, obtaining a partitioning matrix for each set, *P^A^_Train_* and *P^B^_Test_* (if *N* is the total number of neurons, the partitioning matrix *P* is a binary *N*x*N* matrix, where matrix position *P_i,j_* is set to 1 if neuron *i* and neuron *j* belong to the same cluster, and set to 0 otherwise). We then constructed an additional partitioning matrix *P^A->^ ^B^* by using the centroids of clusters in *P^A^_Train_* to perform clustering on set *B*: we assigned neuron *k* to cluster *j* in *P^A->^ ^B^* if ***RelContrib^B^****_k_* was closest to the centroid of cluster *j* across all cluster centroids in set *A*. To quantify generalizability of clustering results, we calculated the sample correlation coefficient between the upper triangular block of matrix *P^B^_Test_* and the upper triangular block of matrix *P^A->^ ^B^*. We repeated this procedure using *B* as train set and *A* as test set, and varied *thresh_cut_* along a range of values. Through this procedure, we determined that the optimal *thresh_cut_* was 2.2 for first-session data, and 2 for expert-session data. For these values of *thresh_cut_*, generalizability of first-session data was 0.79 +/- 0.006 (for shuffled test partitionings it was 0.002 +/- 0.032) and of expert-session data was 0.69 +/- 0.09 (for shuffled test partitionings it was 0.003 +/- 0.030).

To visualize the organization of the relative contribution vectors of all neurons, and perform a qualitative validation of the obtained clusters, we used multidimensional scaling (MDS). This nonlinear technique maps the collection of neuron’s relative contribution vectors {***RelContrib****_i_*}*_i_* from their original eight-dimensional space (corresponding to the 8 task variables) to a bidimensional projection that best preserves the original distances between vectors {𝑝𝑑𝑖𝑠𝑡_𝑖,𝑗_}_𝑖,𝑗_. Neurons with similar relative contribution vectors occupy similar regions of the bidimensional MDS plane, allowing visualization of the global structure of similarities of {***RelContrib****_i_*}*_i_*. We applied the non-classical (non-metric) MDS algorithm on the collection of 𝑝𝑑𝑖𝑠𝑡_𝑖,𝑗_values, calculated with the euclidean distance metric, with Kruskal’s normalized stress-1 goodness-of-fit criterion. In Fig. 3g, neurons in the MDS plane were colored according to the cluster to which they belong, as determined by our hierarchical clustering approach (same color-code used in Extended data Fig. 7).

## Acknowledgments

We thank members of the Marin-Burgin lab, the Refojo lab and Muraro lab for insightful discussions. We thank Emilio Kropff and Massimo Scanziani for valuable comments on the manuscript. We thank Karel Svoboda and Hidehiko Inagaki for the initial instruction for the in vivo recordings; also to Miho Inagaki for the initial assistance in behavioral training of head-fixed mice; and to the Janelia HHMI Institute for hosting NF. We acknowledge International Development Research Centre IDRC108878 (AMB), Argentine Agency for the Promotion of Science and Technology PICT2018-0880 (AMB), PICT2020-00360 (AMB), PICT 2020-1536 (NF), PICT 2016-2758 (NF), PICT 2017-4023 (SAR), CONICET PIP 2787 (NF and SAR) and FOCEM-Mercosur COF 03/11. Fulbright Foundation and 2015-BECar-Argentine Presidential Fellowship in Science and Technology for financial support to NF. We thank Ivan Refojo for the help with the 3D printer. Finally, we thank our National Football team for giving us the joy of being World Champions.

## Author contributions

The project was originally conceptualized by A.M-B, N.F and S.A.R. The behavioural paradigm was developed and designed by N.F., S.A.R. and A.M-B. Task-related hardware was developed and constructed by S.A.R. and N.F. Task-related software was developed and implemented by S.A.R. Animal training, behavior data collection and behavioural data analysis was performed by N.F., S.A.R., M.A-D, L.S. Neural recordings were performed by N.F., S.A.R., L.S., A.M-B. Neural data analysis was performed by N.F. and S.A.R. The manuscript was written by N.F., S.A.R. and A.M-B. and edited and reviewed by all authors.

**Supplementary Information** is available for this paper.

## Data availability

Data will be made available upon reasonable request to the corresponding authors.

